# Neoantigens drive adoptively transferred CD8 T cells to long-lived effectors mediated by lymph node trafficking

**DOI:** 10.1101/2025.04.18.649565

**Authors:** Megen C. Wittling, Amalia M. Rivera Reyes, Megan M. Wyatt, Aubrey S. Smith, Anna C. Cole, Guillermo O. Rangel Rivera, Michael B. Ware, Ayana T. Ruffin, Soundharya Kumaresan, Frances J. Bennett, Gregory B. Lesinski, Chrystal M. Paulos, Hannah M. Knochelmann

## Abstract

Adoptive transfer of neoantigen-reactive T lymphocytes mediates potent responses against solid tumors in patients while often limiting toxicity to normal tissues; however, the mechanisms governing the trafficking, activation, and superior antitumor function of infused T cells remain unknown. Using a clinically relevant TCR-transgenic T cell therapy model, we examined CD8 T cell responses to melanoma expressing either wild-type or mutated antigen. Neoantigen expression conferred robust tumor regression, durable cures, and long-term protective immunity. Mechanistically, T cells reacting to neoantigen exhibited enhanced cytokine and chemokine production, heightened effector function, and sustained persistence within the blood, tumor, and lymph nodes. Notably, trafficking through secondary lymphoid organs was necessary for T cell persistence and antitumor efficacy. These findings highlight the critical role of T cell trafficking to the lymph nodes in shaping neoantigen-specific antitumor responses and offers insight for improving adoptive cellular therapies.

## Introduction

Adoptive T cell transfer (ACT) therapy is an effective treatment for patients with solid tumors, with recent FDA approvals of lifileucel (tumor infiltrating lymphocyte (TIL) therapy for anti-PD1-resistant metastatic melanoma) and afamitresgene autoleucel (TCR-based therapy for patients with metastatic synovial sarcoma)(1,2). As additional cellular therapies emerge(3), a key determinant of their efficacy is the tumor antigen(s) targeted by transferred T cells(4-6). T cell receptors (TCRs) recognize peptides presented on MHC molecules by antigen-presenting cells (APCs) or tumors, which can originate from either nonmutated self-proteins (wild-type) or neoantigens. Neoantigens—novel peptides generated by mutations—are attractive targets because they are largely absent from normal tissue, reducing off-target toxicity(7,8).

Multiple immunotherapies demonstrate enhanced effectiveness when they induce or harness T cell responses against neoantigens. In patients receiving immune checkpoint blockade (ICB) therapy, neoantigen-specific T cells are enriched(5,9-11) and correlate with improved outcomes(12,13). Similarly, ACT therapies targeting neoantigens have induced tumor regression in patients with difficult to treat cancers, including pancreatic cancer, colon cancer, and cholangiocarcinoma(14-17). Neoantigen-based tumor vaccines are also being explored in combination with ICB and ACT to further amplify antitumor responses(18,19). Yet how the mutational status of a tumor antigen—neoantigen versus non-mutated self-antigen—affects the fate and function of T cell therapy products remains unclear.

To address this, we used the B16F10 or B16KVP melanoma tumor models, engineered to express either the wild-type non-mutated or neoantigen forms of gp100(20) respectively, in order to track how an identical T cell product traffics, persists, and mediates anti-melanoma activity in these two different contexts. Prior studies suggest that pmel-1 T cell therapy exhibits enhanced activity against neoantigen expressing tumors(20), but the mechanisms underlying this advantage remain undefined. Here, we dissect how T cell phenotype, cytokine profile, and gene expression change when adoptively transferred T cells engage with wild-type versus neoantigen-expressing tumors. This study provides a rare opportunity to define tumor-specific T cell dynamics across tissues in a controlled setting. Our findings reveal unexpected insights into the superior efficacy of neoantigen-specific T cells – illuminating the kinetics of effector differentiation in the context of mutated versus nonmutated tumor.

## Materials and Methods

### Mice

Both pmel-1 (B6.Cg-*Thy1*^*a*^/Cy Tg(TcraTcrb)8Rest/J) and C57BL/6 mice were purchased from Jackson Laboratories and housed in the in-house animal facilities at Emory University. All mice were used in accordance with the Emory University Institutional Animal Care and Use Committee guidelines. Pmel-1 (B6.Cg-*Thy1*^*a*^/Cy Tg(TcraTcrb)8Rest/J) are bred in house at Emory University. C57BL/6J mice were purchased from Jackson laboratories at 6-8 weeks. Female C57BL/6J mice were used for all experiments. C57BL/6J mice were randomized prior to treatment, and mice with either no detectable or intraperitoneal tumors were excluded from studies. Tumors for mice were measured a minimum of 2x per week with increasing measurements as tumor burden scores increased. They were measured length by width using handheld calipers by a laboratory member blinded to the treatment group. Mice were euthanized either when they reached endpoint or when scheduled for biodistribution/tissue collection.

### Media for Tumor and Cell Culture

Complete Media is made by combining RPMI 1640 w/ L-glutamine (1L), 10% HI FBS (100mL), 1% Pen/Strep (10mL), 1% Non-essential Amino Acids (10mL), 1% Sodium Pyruvate (10mL), 0.1% HEPES (1mL), and 0.1% 2-Mercaptoethanol (1mL). After combining, the media is sterile filtered and stored at 4 degrees Celsius for up to one month.

### Tumor Cell Lines

B16F10 is available from ATCC (CRL-6475). B16KVP tumor cell lines are a gift from Nicholas Restifo. They are cultured in complete media. When splitting, 0.05% Trypsin is used.

### T Cell Culture

Pmel-1 spleens were processed over a 70 μm filter. Red blood cell lysis was then performed using 1x RBC lysis buffer added for 5 minutes. Cells were then resuspended in media at 1e6 cells/mL. For the expanded pmel-1 cell experiment, cells were plated with 2mL of complete media in a 24-well plate and activated with 1□µM of human gp100 (hgp100) peptide along with 100 IU/mL of IL-2 for 7 days. For all other experiments, cells were not activated.

### Naïve Cell Sort

Pmel-1 T cells were harvested as described above, and naïve T cells were then collected using the EasySep Mouse Naïve CD8+ T Cell Isolation Kit per manufacturer’s instructions. Prior to infusion into mice, T cells were assayed for their CD44-CD62L+ phenotype and pmel-1 purity via flow cytometry.

### Tumor Injection

B16F10 or B16KVP tumor cells were harvested, washed twice in sterile PBS, and then resuspended at a concentration of 500,000 cells/200uL. 200uL was then injected subcutaneously onto the flank of mice. Tumors were allowed to established for one week prior to adoptive cell transfer. For rechallenge experiments, B16KVP tumor cells were washed and resuspended in sterile PBS, and 200μL containing 500,000 B16KVP cells was subcutaneously injected onto the flank of mice.

### Lymphodepletion

Mice were subjected to 4-5Gy of total body irradiation (TBI) one day prior to adoptive T cell transfer.

### Adoptive Cell Therapy

1e6 pmel-1 T cells (either sorted naïve cells or expanded cells as described above) were resuspended in sterile PBS and transferred via tail vein injection into C57BL/6 mice.

### FTY720 Administration

FTY720 (Cayman Chemicals #10006292) was administered at a dose of 1mg/kg for each mouse. The drug was resuspended in 100uL of sterile water for each mouse and injected intraperitoneally daily for a total of 10 days beginning the day of adoptive T cell therapy.

### Blood and Tissue Collection

#### Peripheral Blood

Blood from the mandibular vein was collected into EDTA coated tubes. Tubes were spun at 1800 rpm for 10 minutes, after which serum was collected and stored at -80 degrees Celsius for future cytokine multiplex analysis. Blood was then lysed using RBC lysis buffer for 5 minutes. Cells were then resuspended in FACS Buffer and assayed via flow cytometry.

#### Tumor

B16F10 or B16KVP tumors were excised into a plate containing complete media after which they were then minced using a sterile scalpel and processed over a 70 μm filter. Cells were then washed twice in sterile PBS and then assayed via flow cytometry.

#### Lymph Nodes

The draining lymph node (inguinal) was excised from mice and plated in media, after which they were processed over a 70 μm filter. Cells were then washed twice in sterile PBS and then assayed via flow cytometry.

#### Spleen

Spleens were excised from mice and plated in media, after which they were processed over a 70 μm filter. Cells were centrifuged at 1400 rpm for 5 minutes. RBC lysis was then performed for 5 minutes using 1X RBC lysis buffer. Media was added and cells centrifuged at 1400 rpm for 5 minutes. Cells were then washed twice by centrifugation in sterile PBS and then assayed via flow cytometry.

### Flow cytometry

Flow cytometry was performed on the 5 laser Cytek Aurora system and subsequently analyzed using FlowJo software. For viability staining, cells were resuspended in PBS and fixable viability dye for 15 minutes at room temperature. Cells were then washed twice and resuspended in FACS buffer (PBS+2%□FBS) with a 1:500 dilution of extracellular antibodies listed in **Supplemental Table 2** for 20 minutes at room temperature. For intracellular stains, the FoxP3 transcription factor kit was used (ThermoFisher 00-5523-00). Cells were first resuspended in fixation/permeabilization buffer for 15 minutes and subsequently washed in 1x permeabilization buffer. They were then stained in a 1:100 dilution of intracellular antibodies in Permeabilization Buffer for 30 minutes at room temperature (**Supplemental Table 2**). Cells were then washed with FACS buffer prior to running on the Aurora.

### Cytokine Multiplex Analysis

Serum was collected from mice 5 days after adoptive T cell therapy of naïve pmel-1 T cells and was stored at -80 degrees Celsius until ready for analysis. There were seven to twelve mice per group. A 44-plex cytokine array was performed on the serum samples by Eve Technologies according to manufacturer’s protocol (Mouse Cytokine/Chemokine 44-Plex Discovery Assay).

### Nanostring

Tumor tissue was collected 5 days after adoptive T cell therapy, placed in RNAlater solution and stored at 4 degrees Celsius. There were three tumor samples collected per group. The samples were then given to the Emory Integrated Genomics Core for RNA extraction and quality control. Nanostring was then conducted using the Nanostring PanCancer IO 360 Panel. For Nanostring analysis, RCC files and the RLF file for the IO360 Panel were uploaded into nSolver (Bruker). Background thresholding was estimated from Negative control counts using the geometric mean of negative controls. For normalization, positive control normalization using the geometric mean and CodeSet Normalization using the Standard and geometric means was performed. Fold change estimation compared the B16KVP tumored mice to the B16F10 tumored mice that received pmel-1 ACT, and a false discovery rate (FDR) was calculated. A list of all significant genes is provided in **Supplemental Table 1**. For volcano plots and graphs depicting pathways and differentially regulated genes, a p-value of 0.05 was used as the cutoff to determine genes of significance. Volcano plots depict a log2 fold change cutoff of ± 0.5 and were made using R and RStudio.

### Pathway Analysis

GO Enrichment Analysis was performed using the Gene Ontology Resource. The significant upregulated genes were inputted into the PANTHER GO Enrichment tool that connects to up-to-date GO annotations. GO Biological Processes for Mus musculus were utilized for analysis and significantly upregulated pathways visualized as the -log10 of the False Discovery Rate (FDR).

### Survival Analysis

The top 10 genes that were significantly upregulated in neoantigen tumors day 5 post-ACT were evaluated for their impact on survival in the TCGA SKCM cohort. Overall Survival with a Median Group Cutoff was used. Hazards Ratio was calculated based on the PH model. 95% confidence interval is displayed as a dotted line. Graphs made using GEPIA.

### Statistical Analysis

All measurements displayed were taken from distinct samples with no repeated measures of the same sample. Graphs and statistics were performed using the GraphPad Prism software. Survival analyses were performed using the Log-rank (Mantel-Cox) test. For comparisons between 2 treatment groups, a two-sided Mann-Whitney U test was performed as indicated. For comparisons between >2 groups, a Kruskal-Wallis test with Dunn’s multiple comparisons test was performed. Nonparametric tests were utilized for all statistical comparisons due to sample size and violation of normality assumptions.

### Data and materials availability

Nanostring data generated from this study can be located at GSE285952. All other data is available upon reasonable request.

## Results

### Neoantigen recognition enhances the efficacy of pmel-1 T cells against melanoma

We tested whether a CD8 T cell therapy product would exhibit distinct antitumor activity in mice bearing melanoma expressing either a nonmutated or mutated antigen. To do this, we used the pmel-1 transgenic T cell system, in which all CD8 T cells express a TCR specific for the gp100 antigen. This model allowed us to directly compare responses against B16F10 melanoma (expressing wild-type gp100) and B16F10-KVP melanoma (expressing a neoantigen form of human gp100 with the KVP mutation)(20). Both tumors have similar growth kinetics in mice, providing a comparable system to evaluate antigen-specific T cell responses. Since neoantigens arise from mutations in endogenous tumor antigens, this model enables the study of T cell responses to neoantigens in otherwise genetically identical melanoma cell lines.

As outlined in **Figure 1A**, we transferred either naïve or IL-2–expanded anti-melanoma pmel-1 CD8 T cells into mice bearing B16F10 or B16KVP tumors. IL-2–expanded T cells reflect a clinically relevant approach for expansion of adoptive cell transfer (ACT) products(21). We expected T cells targeting the neoantigen to mount a stronger antitumor response than those recognizing wild-type antigen. While a transient regression of B16KVP neoantigen tumors was seen, the IL-2-expanded pmel-1 T cells failed to provide durable tumor control (**Figure 1B**) against both tumor types.

**Figure 1:**
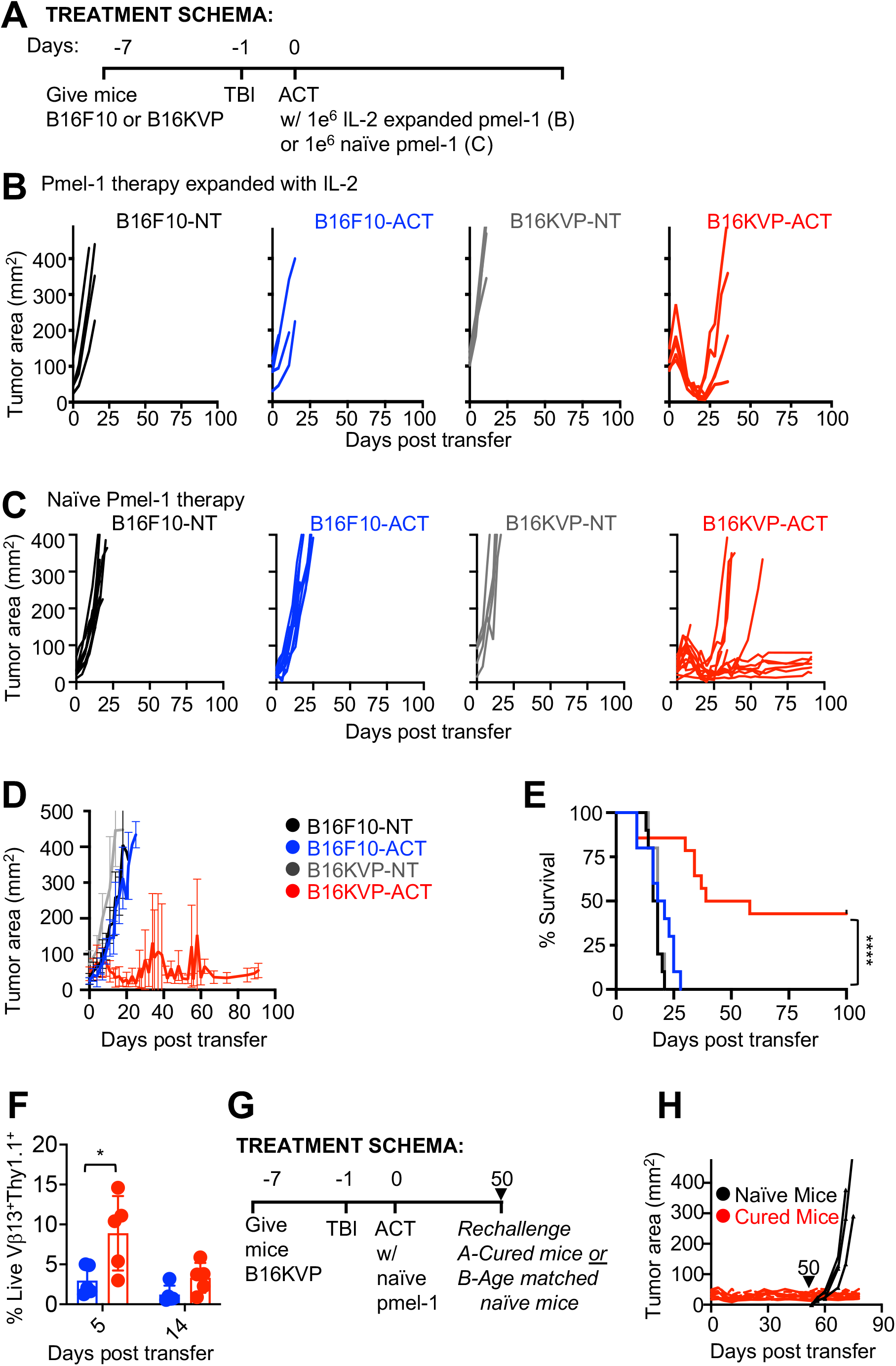
Pmel-1 CD8 T Cells effectively regress neoantigen but not wild-type melanoma tumors. **(A)** Experimental setup of adoptive T cell therapy into neoantigen versus wildtype tumors. **(B)** Individual tumor curves of mice given IL-2 expanded T cell therapy to neoantigen or wildtype tumors. (n=4 mice/group) **(C)** Individual tumor curves of mice given naïve pmel-1 CD8 T cell therapy to neoantigen or wildtype tumors (n=10-14 mice/group). **(D)** Overlay of tumor curves from all groups displayed in Figure 1C. **(E)** Survival of mice with B16F10 or KVP tumors ± naïve pmel-1 therapy. (**** denotes significant p<0.0001). p-value determined via Log-rank test. **(F)** Engraftment of adoptively transferred naïve pmel-1 cells in the blood on day 5 and day 14 post-transfer. (* denotes significant p < 0.05; p= 0.03). P-value determined via Mann-Whitney U test. (n=5/group) **(G)** Schematic of tumor rechallenge experiment. **(H)** Tumor curves of previously cured mice (red) or treatment-naïve mice (black) rechallenged with B16KVP. (n=4 or 10 mice/group).

Given the established role of naïve and stem-like T cells in mediating effective antitumor responses,(22-25) we hypothesized that naïve pmel-1 CD8 T cells might be more effective than the more differentiated IL-2 expanded T cells. To test this, naïve pmel-1 CD8 T cells were enriched from the spleens of pmel-1 transgenic mice using magnetic bead sorting and assessed for their response to mutated B16KVP or nonmutated B16F10 tumors. Flow cytometry confirmed that the majority of isolated pmel-1 T cells were naïve (**94%:** CD44^-^CD62L^+^; **Supplemental Figure 1**).

Naïve pmel-1 T cells failed to control B16F10 wild-type tumors but mounted a robust response against B16KVP neoantigen-expressing tumors, leading to durable tumor regression in >50% of mice and improved long-term survival (**Figure 1C-E**). Remarkably, this effect occurred without exogenous IL-2 or vaccine co-administration, both typically required to drive responses in the B16F10 model(25-27). Additionally, transferred pmel-1 CD8 T cells engrafted at a significantly higher percentage in mice with neoantigen-expressing melanoma tumors at day five post-transfer (p = 0.03), with a trend towards increased persistence by day 14 (**Figure 1F**).

To further investigate the durability and memory potential of naïve pmel-1 T cell-mediated tumor immunity, we next examined whether mice ablated of their primary B16KVP tumors could mount protective immunity upon tumor rechallenge (**Figure 1G**). Establishing long-term immunological memory is a critical hallmark of effective T cell–based therapies, ensuring lasting immunity against tumor recurrence. Notably, mice that had been cured of B16KVP tumors through naïve pmel-1 adoptive cell therapy (ACT) were fully protected upon rechallenge, while age-matched, treatment-naïve control mice developed aggressive tumors and succumbed to disease (**Figure 1H**). These findings suggest that naïve pmel-1 T cells not only mediate robust initial tumor clearance but also provide lasting immunity against neoantigen-expressing tumors. Yet the mechanisms by which naïve pmel-1 T cells drive immunological memory to neoantigen-expressing tumors remains poorly understood.

### Signatures of cytokine signaling, immune cell recruitment, and antigen presentation are increased after therapy in B16KVP mice

To understand the mechanisms underlying the robust responses observed in the neoantigen-expressing tumors, we examined the molecular and immunological landscape of tumors following naïve pmel-1 T cell infusion. Despite both tumors receiving the same initial T cell product, B16KVP tumors regressed while B16F10 tumors progressed (**Figure 1**), prompting us to investigate the immune differences mediating these outcomes.

RNA transcript profiling of tumor tissues at day five post-T cell transfer revealed over 200 differentially expressed genes in the B16KVP tumors relative to B16F10 tumors (**Figure 2A, Supplemental Table 1**). Among the top 50 upregulated genes in B16KVP tumors were those associated with cytokine signaling (*Irf8, Cxcl9, Cxcl10*) and antigen presentation (*H2-D1, H2-M3, H2-K1*) (**Figure 2B**). Gene ontology (GO) enrichment analysis further identified biological processes associated with cell activation, cytokine responses, and leukocyte differentiation to be higher in B16KVP tumors (**Figure 2C**)(28-30). These findings indicate that, even at early time points (day five post-ACT), neoantigen-expressing tumors permit a more immunogenic microenvironment than their wild-type counterparts.

**Figure 2:**
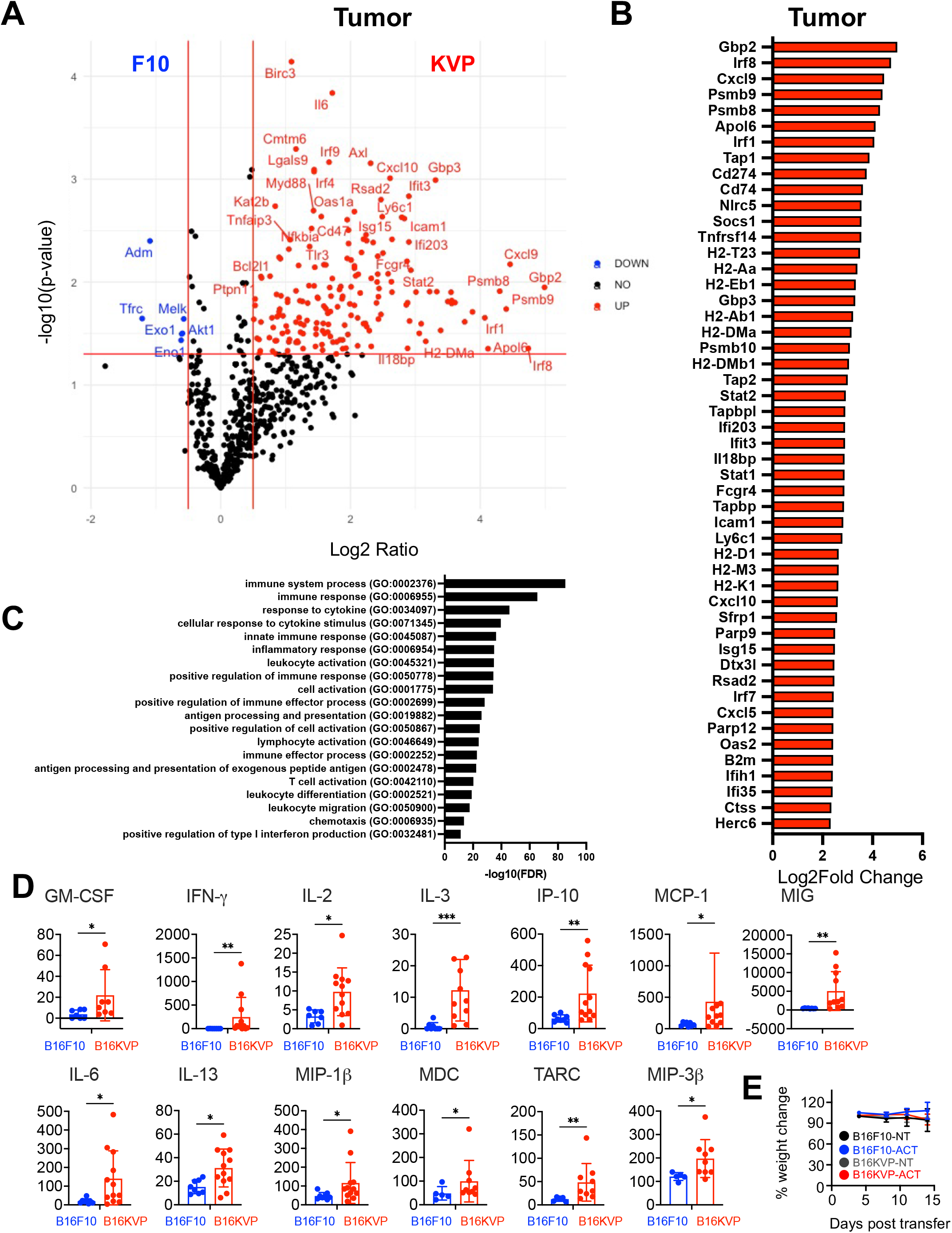
Increased inflammatory cytokine signaling and effector processes are found in the tumor of neoantigen tumors as compared to wild-type. **(A)** Volcano plot depicting differentially expressed genes in the tumor between mice with neoantigen versus wild-type tumors and given naïve CD8 pmel-1 T cells (n=3 mice/group). Red lines depict p-value of 0.05 and log2 fold change of ± 0.05. **(B)** Top 50 differential genes upregulated in neoantigen-bearing mice that are statistically significant (p<0.05). **(C)** Pathway analysis of significantly upregulated genes enriched in the neoantigen tumored mice. GO enrichment analysis, biological pathways used. **(D)** Serum collected from mice with B16KVP or B16F10 tumors five days after receiving adoptive transfer of naïve pmel-1 T cells (n=7-12 mice/group). Serum cytokines were profiled using Eve Technologies mouse cytokine/chemokine discovery assay array. Mann-Whitney U test performed. **(E)** Percentage of weight change for mice with B16F10 or B16KVP tumors +/-pmel-1 cellular therapy (n=5 mice/group). Weight measured for two weeks post-transfer.

To evaluate the clinical relevance of these findings, we examined the top ten most upregulated genes in B16KVP tumors (*Gbp2, Irf8, Cxcl9, Psmb9, Psmb8, Apol6, Irf1, Tap1, Cd274, Cd74*) and evaluated their correlation with patient outcomes using publicly available TCGA data via GEPIA(31). Notably, all ten genes were positively associated with improved survival in patients with skin cutaneous melanoma (SKCM); **Supplemental Figure 2**. These data suggest that genes enriched in neoantigen-bearing tumors may have prognostic significance in human melanoma.

### Neoantigen-specific tumors induce a systemic pro-inflammatory cytokine signature

Given the enrichment of transcripts associate with cytokine signaling pathways in B16KVP tumors, we sought to explore which cytokines may be potentiating the antitumor responses observed. To test this, we collected serum from mice bearing B16KVP or B16F10 tumors on day five post-transfer of naïve pmel-1 CD8 T cells. Over 40 cytokines were profiled, and a broad array of cytokines were elevated in the serum of mice with neoantigen tumors compared to those with wild-type tumors (**Figure 2D, Supplemental Figure 3**).

**Figure 3:**
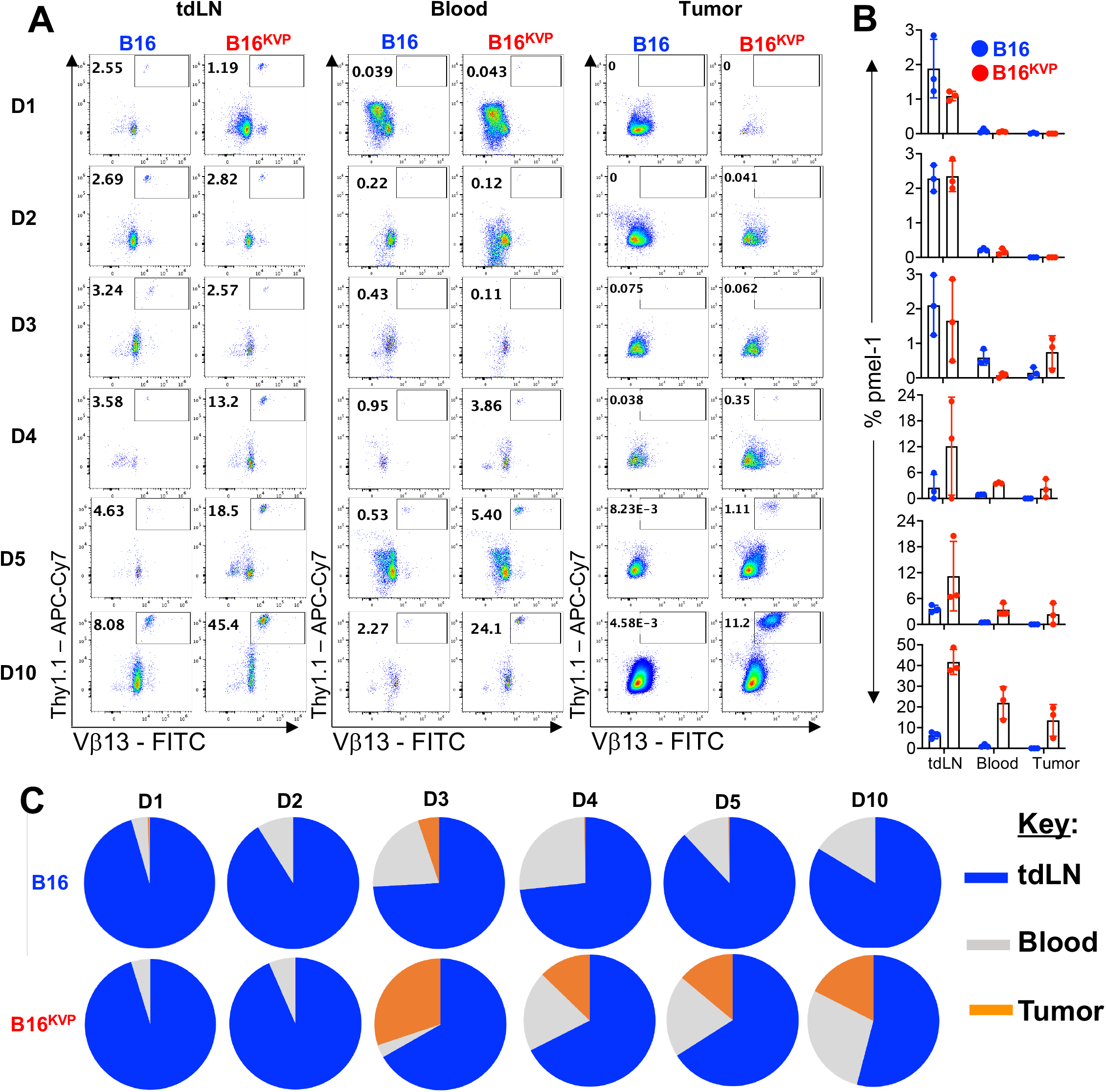
Evaluating the kinetics of adoptively transferred naïve CD8 T Cells in the context of neoantigen or non-mutated tumors. **(A)** Representative flow plots of percentage of transferred pmel-1 T cells (Vb13+ Thy1.1+) in either B16F10 or B16KVP tumored mice at various time points in the tdLN, blood, and tumor. **(B)** Bar graph representation of percentage of pmel-1 cells in the tdLN, blood, and tumor at each time point (n=3 mice/group). **(C)** Pie charts depicting proportion of transferred cells in the tdLN, blood, and tumor at each time point.

Notably, interferon-gamma (IFN-g), a key pro-inflammatory cytokine, exhibited a >300-fold increase in B16KVP as compared to B16F10 tumors (**Figure 2D**). Additionally, cytokines known for immune cell recruitment — including IP-10 (CXCL10), MCP-1 (CCL2), MIG (CXCL9), MIP-1b (CCL4), MDC, TARC (CCL17), and MIP-3b (CCL19) —were significantly increased in the mice bearing B16KVP tumors (**Figure 2D**)(32-34). Interleukins such as IL-2, IL-3, and IL-6 were also significantly increased and may play a key role in mediating immunity to neoantigen tumors (**Figure 2D**). It is also important to note that while many immunomodulatory cytokines were increased in B16KVP-bearing mice, mice displayed little toxicity as measured by nominal weight changes over time post-ACT therapy (**Figure 2E**).

### Adoptively transferred T cells rapidly traffic to the lymph node and preferentially expand in neoantigen-expressing tumors

As robust genetic changes were detected in B16KVP tumors after ACT, we hypothesized that the transferred T cells were primarily trafficking to the neoantigen-rich melanoma to exert their effector functions. To evaluate the migration pattern of infused pmel-1 T cells, a kinetic analysis of T cell trafficking and persistence was performed across multiple tissues over time.

Within the first two days post-transfer, pmel-1 CD8 T cells were detectable at low frequencies in lymph nodes of mice bearing either B16F10 (wild-type) or B16KVP (neoantigen-expressing) tumors (**Figure 3A**) but were not detected in the blood or tumor. However, within three to four days post-transfer, a divergence emerged. Pmel-1 T cells began infiltrating B16KVP neoantigen tumors but were nearly absent in wild-type B16F10 tumors of mice (**Figure 3A,B**). In the B16KVP group, T cell frequencies within the tumor steadily increased overtime (four, five, or ten days post-transfer), indicating their sustained expansion and accumulation. In contrast, few (>1%) pmel-1 T cells were detectable in wild-type B16F10 tumors across all time points (**Figure 3A, 3B**).

A comprehensive comparison of the distribution of transferred cells across the lymph node, blood, and tumor highlights this pattern (**Figure 3C**). Initially, pmel-1 T cells were present at similar proportions in mice bearing either malignancy. However, by day three onward, only mice with neoantigen-expressing tumors maintained a substantial population of pmel-1 T cells within the blood and the tumor. This progressive enrichment in B16KVP tumors, coupled with their sustained presence across all tissues from ∼four to ten days post-infusion suggests that neoantigen recognition allows sufficient activation and expansion signals for T cells to thrive in multiple host environments.

### Adoptively transferred pmel-1 CD8 T cells proliferate and acquire effector-like phenotypes in the neoantigen tumor-draining lymph nodes

To examine the impact of neoantigen tumor expression on T cell activation and differentiation for ACT therapy, naïve pmel-1 CD8+ T cells were transferred into mice bearing either neoantigen-expressing B16KVP tumors or wild-type B16F10 tumors. Prior to ACT, baseline characterization confirmed that enriched naïve pmel-1 cells expressed low levels of activation markers (CD44, ICOS, Tim-3, CD69) and maintained a CD62L+CD44?TCF1+PD-1^-^ profile compared to bulk populations (**Supplemental Figure 4**).

**Figure 4:**
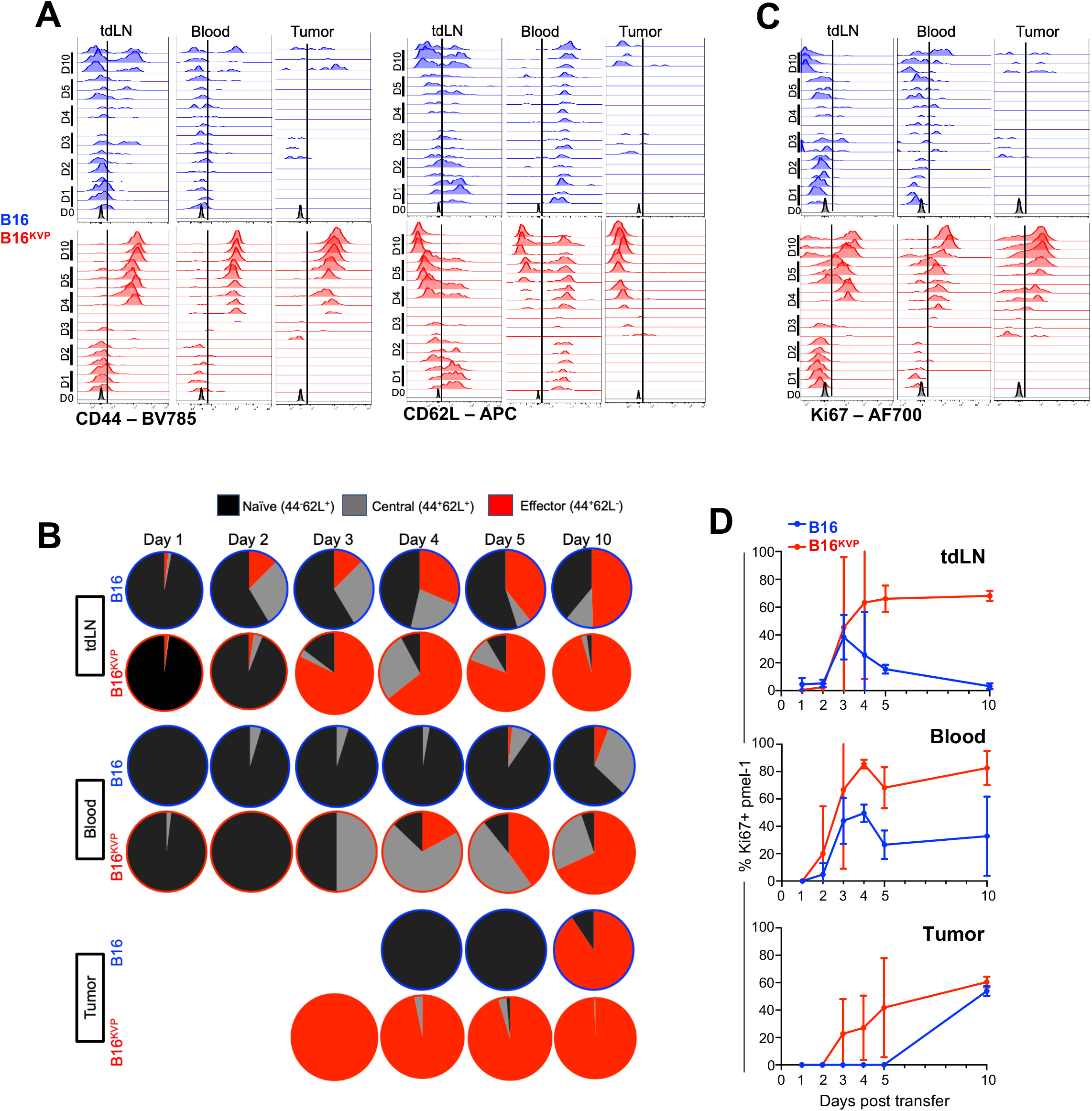
Assessing proliferation and phenotype of adoptively transferred cells in r wild-type context. **(A)** Histograms displaying CD44 and CD62L expression by the transferred pmel-1 T cells at various time points in the tdLN, blood, and tumor in either neoantigen or wild-type tumor bearing mice. **(B)** Pie charts depicting percentage of transferred pmel-1 cells that are naïve (CD62L+CD44-), central memory (CD62L+CD44+), or effectors (CD62L-CD44+). **(C)** Histograms displaying Ki-67 expression in the transferred pmel-1 T cells at various time points in the tdLN, blood, and tumor in either neoantigen or wild-type tumor bearing mice. **(D)** Graphical representation of Ki-67 expression over time in the tdLN, blood, and tumor (n=3 mice/group).

Kinetic analysis revealed that, despite their naïve origin, pmel-1 cells in B16KVP-bearing mice rapidly acquired an effector phenotype (determined by CD44 and CD62L expression via flow cytometry). By three to four days post–transfer, these cells upregulated CD44 – a hallmark of activation and differentiation – in the blood, tumor, and tdLN (**Figure 4A**). Concurrently, CD62L expression was progressively downregulated in the tdLN, consistent with a transition from a naïve or central memory state to an effector phenotype (**Figure 4A and 4B**). In contrast, T cells from mice bearing B16F10 tumors exhibited significantly attenuated CD44 upregulation and maintained higher levels of CD62L.

We additionally evaluated proliferation by measuring expression of Ki-67 in pmel-1 cells by flow cytometry. In mice bearing B16KVP tumors, pmel-1 T cells exhibited sustained and progressively increased Ki-67 expression across all compartments compared to mice bearing wild-type B16F10 tumors. (**Figures 4C and 4D**).

Collectively, these findings reveal that adoptively transferred naïve antigen-specific T cells differentiate into effector-like cells in the presence of neoantigen-expressing tumors as well as proliferate at much higher levels. This effector transition is first seen in the draining lymph node and is sustained as T cells traffic to and expand within the neoantigen-expressing tumors, ultimately supporting potent and durable antitumor immunity. These data indicate that neoantigen expression not only enhances the proliferative capacity of adoptively transferred T cells but also accelerates their effector differentiation.

### Neoantigen-expressing tumors induce activation and effector differentiation of adoptively transferred T cells

To assess how neoantigen-expressing tumors modulate the activation, exhaustion, and stemness profiles of adoptively transferred naïve pmel-1 T cells, we performed an extensive phenotypic analysis on day ten post–ACT. High frequencies of pmel-1 T cells were detected in all examined tissues of B16KVP tumor-bearing mice (**Figure 3A**), enabling detailed profiling.

Notably, pmel-1 T cells expressed elevated PD-1 levels across the lymph nodes, blood, tumor, and spleen, indicating persistent antigen engagement. Strikingly, approximately 33% of tumor-infiltrating pmel-1 T cells co-expressed PD-1 and Tim-3, a phenotype consistent with antigen experience (**Figure 5A**). Furthermore, these cells exhibited increased CD69 expression across all tissues, confirming recent activation (**Figure 5B**).

**Figure 5:**
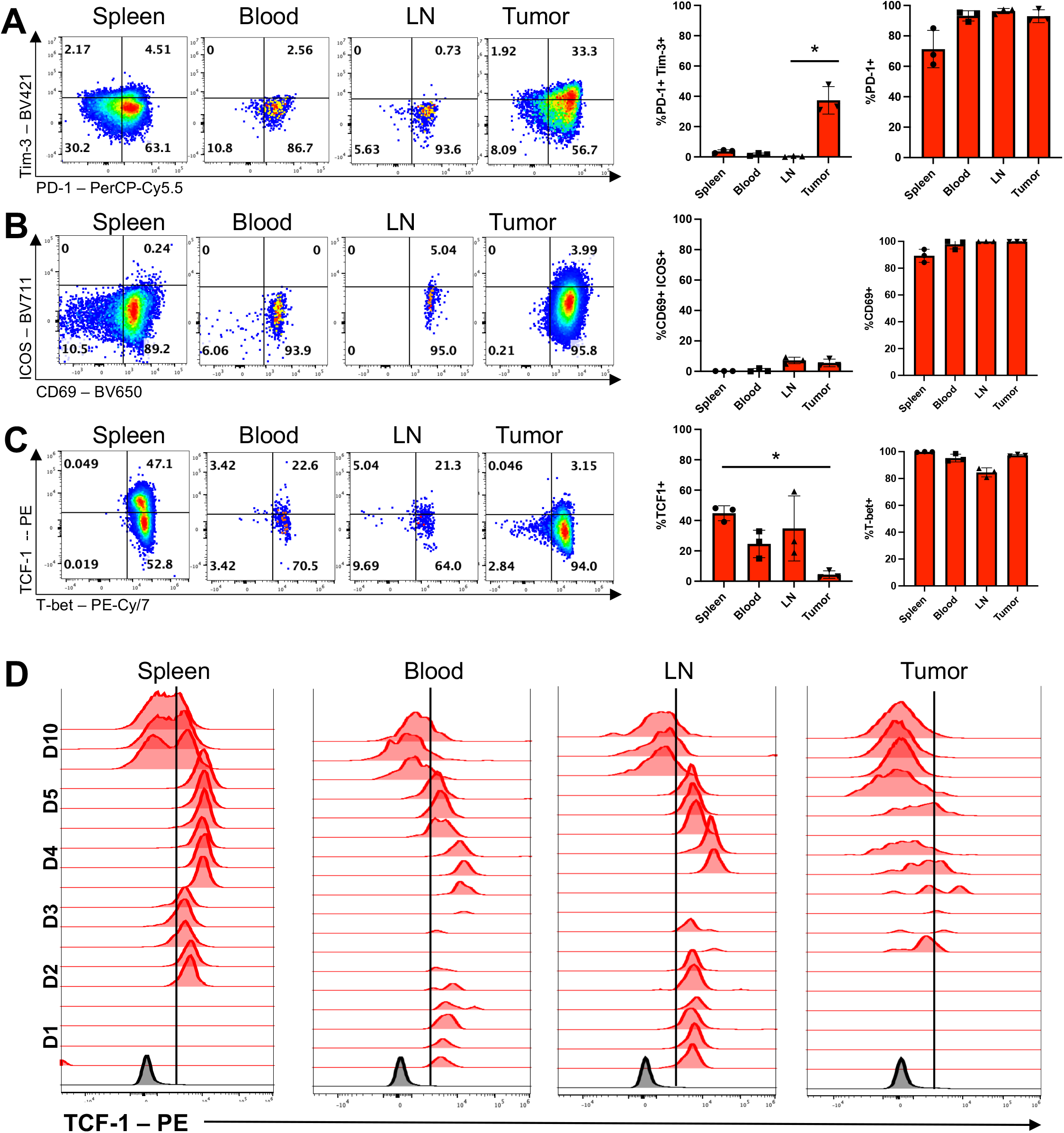
Assessing proliferation and phenotype of adoptively transferred cells in the neoantigen or wild-type context. **(A)** Representative flow plots of PD-1 and Tim-3 expression by the adoptively transferred cells ten days after initial transfer in the blood, tdLN, spleen, and tumor. Accompanying bar graphs depict percentage of cells that are either PD-1 Tim-3 double positive or PD-1 positive (n= mice/group). Kruskal-Wallis test with multiple comparisons performed. **(B)** Representative flow plots of ICOS and CD69 expression by the adoptively transferred cells ten days after initial transfer in the blood, tdLN, spleen, and tumor. Accompanying bar graphs depict percentage of cells that are either CD69 ICOS double positive or CD69 positive (n=3 mice/group). Kruskal-Wallis test with multiple comparisons performed. **(C)** Representative flow plots of TCF1 and T-bet expression by the adoptively transferred cells ten days after initial transfer in the blood, tdLN, spleen, and tumor. Accompanying bar graphs depict percentage of cells that are either TCF1 positive or T-bet positive (n=3 mice/group). Kruskal-Wallis test with multiple comparisons performed. **(D)** Histograms displaying Tcf-1 expression by the transferred pmel-1 T cells at various time points in the spleen, blood, tdLN, and tumor in either neoantigen or wild-type tumor bearing mice.

In addition, robust T-bet expression—a transcription factor associated with cytotoxic effector functions—was detected in all tissues. Conversely, TCF1, a marker of T cell stemness, was markedly reduced in the tumor on day ten while remaining highest in the spleen, suggesting enhanced differentiation at the tumor site (**Figure 5C**). Notably, TCF1 expression was also substantially diminished on pmel-1 T cells within the tdLN, indicating that some infused T cells may progressively lose their stem-like properties and differentiate into effector-like memory cells over time.

To investigate the transition of adoptively transferred naïve pmel-1 T cells from a stem-like to an effector phenotype in mice bearing neoantigen-expressing tumors, we examined TCF1 expression over time across multiple tissues in a B16KVP melanoma model. Within the first three days post-ACT, the majority of pmel-1 T cells in the tumor-draining lymph node (tdLN), blood, and spleen expressed high levels of TCF1, indicative of a stem-like state (**Figure 5D**). However, as time progressed and these cells began infiltrating the tumor, their TCF1 levels decreased – becoming virtually undetectable between days five and ten post-infusion in all organs except for the spleen.

Our findings delineate a three-phase process in this ACT model: (1) rapid, antigen-independent trafficking of naïve pmel-1 T cells to the lymph nodes within one to two days; (2) a robust, neoantigen-dependent expansion in the lymph node, blood, and tumor between three and four days; and (3) sustained, selective infiltration, expansion, and persistence of pmel-1 T cells across multiple tissues by days 5-10 post-transfer. This dynamic response underscores the critical role of neoantigen recognition in licensing T cell persistence across tissues—a feature directly associated with durable tumor clearance.

### Lymph node egress is essential for the antitumor efficacy of transferred T cells

Given that adoptively transferred naïve pmel-1 T cells initially localize to lymphoid tissues before infiltrating tumors, we investigated the role of lymphoid trafficking in mediating therapeutic efficacy. Although previous studies have suggested that T cells acquire effector functions predominantly within the tumor microenvironment(35), we hypothesized that initial priming in secondary lymphoid organs is critical for establishing a durable antitumor response. To test this, we employed FTY720, a sphingosine-1-phosphate receptor 1 (S1PR1) antagonist that effectively blocks lymphocyte egress from lymphoid tissues(36).

Mice bearing B16KVP neoantigen tumors received transferred naïve pmel-1 CD8+ T cells along with daily FTY720 treatment for 10 days, while control mice received no additional intervention (**Figure 6A**). We anticipated that inhibiting lymph node egress would disrupt the formation of a long-lived, primed T cell pool, thereby impairing the antitumor response. To confirm FTY720 activity, we assessed lymphocyte distribution. Comparable levels of host CD3 cells—and similarly, transferred pmel-1 T cells—were detected in the lymph nodes of both FTY720-treated and control mice (**Figures 6B-D,E**). In contrast, FTY720 treatment led to a significant reduction of both host and transferred T cells in the blood, confirming effective blockade of lymphocyte egress (**Figures 6B–E**). Notably, the viability of transferred T cells within the lymph nodes was unaffected by FTY720 (**Supplemental Figure 5**).

**Figure 6:**
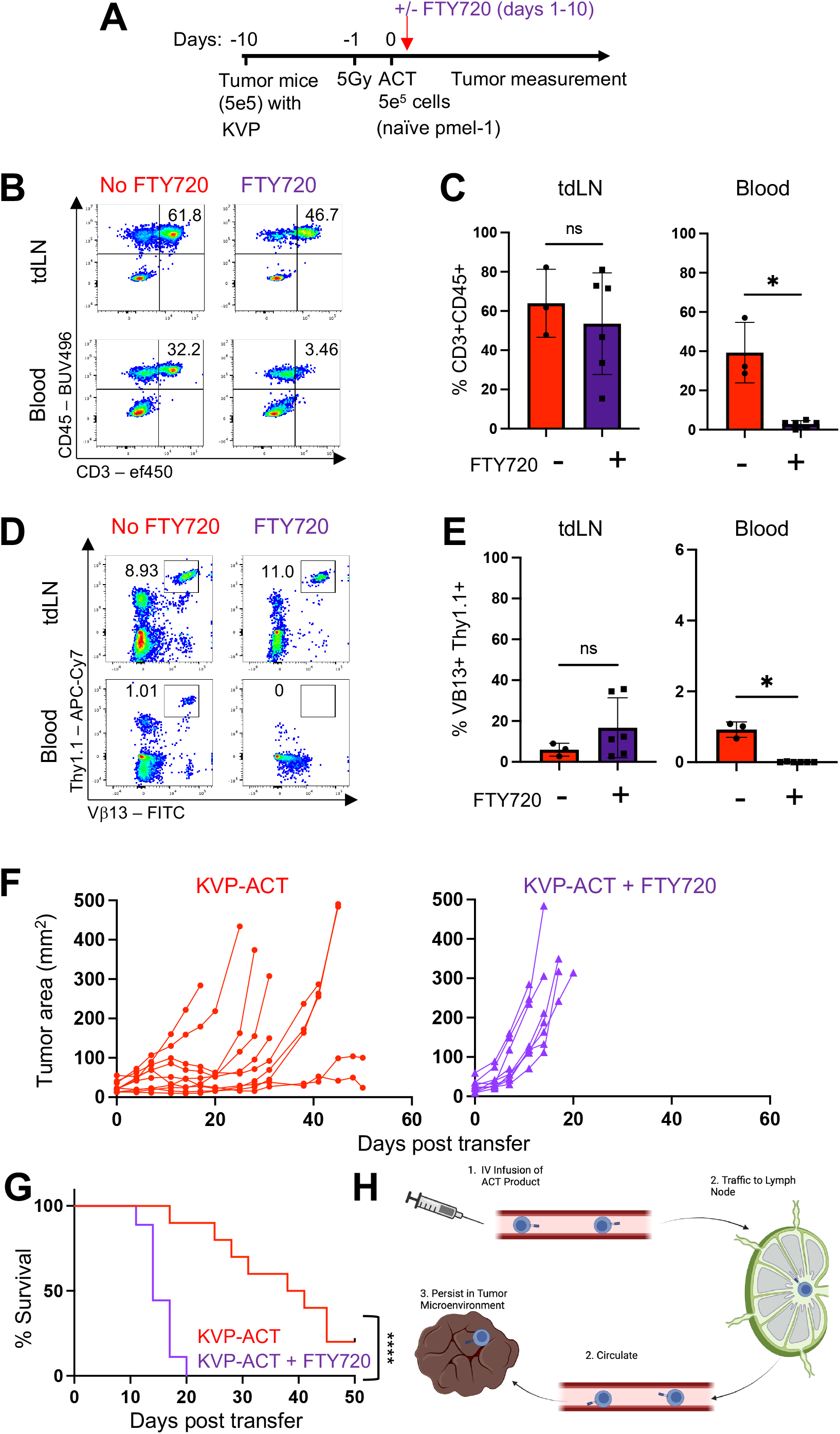
Distinguishing how trafficking to the lymph node impacts therapy response. **(A)** Experimental design schematic shown. **(B)** Lymphocyte presence in the blood or lymph node with or without FTY720 administration as identified via CD45 CD3 dual positivity. **(C)** Bar graphs depicting percentage of lymphocytes in the tdLN and blood with or without FTY720 administration (n=3 or 6 mice/group). **(D)** Representative flow plots of percentage of transferred pmel-1 T cells in the tdLN and blood with or without administration of FTY720. **(E)** Bar graphs depicting percentage of transferred pmel-1 T cells in the tdLN and blood with or without FTY720 administration (n=3 or 6 mice/group). **(F)** Tumor curves for B16KVP-tumored mice given naïve pmel-1 ACT with or without FTY720 administration (n=6 or 10 mice/group). **(G)** Survival curves for B16KVP-tumored mice given naïve pmel-1 ACT with or without FTY720 administration. Log-rank test performed for p-value (<0.0001). **(H)** Schematic depicting proposed trafficking of transferred cells.

Tumor monitoring revealed that blocking lymphocyte egress profoundly impaired therapeutic efficacy. Mice treated with FTY720 exhibited significantly accelerated tumor progression compared to untreated controls (p < 0.0001; **Figures 6F,G**).

These findings underscore that lymphoid egress is essential for the antitumor function of adoptively transferred naïve pmel-1 T cells.

Based on these results, we propose a neoantigen tumor model wherein adoptively transferred T cells first traffic to secondary lymphoid organs to undergo essential priming and early effector differentiation, and subsequently migrate into the tumor to mediate effective, long-term tumor clearance (**Figure 6H**). This work highlights the indispensable role of lymphoid dynamics in driving potent and durable neoantigen-specific antitumor immunity. This collective work is mechanistically visualized in its entirety in **Figure 7**.

**Figure 7:**
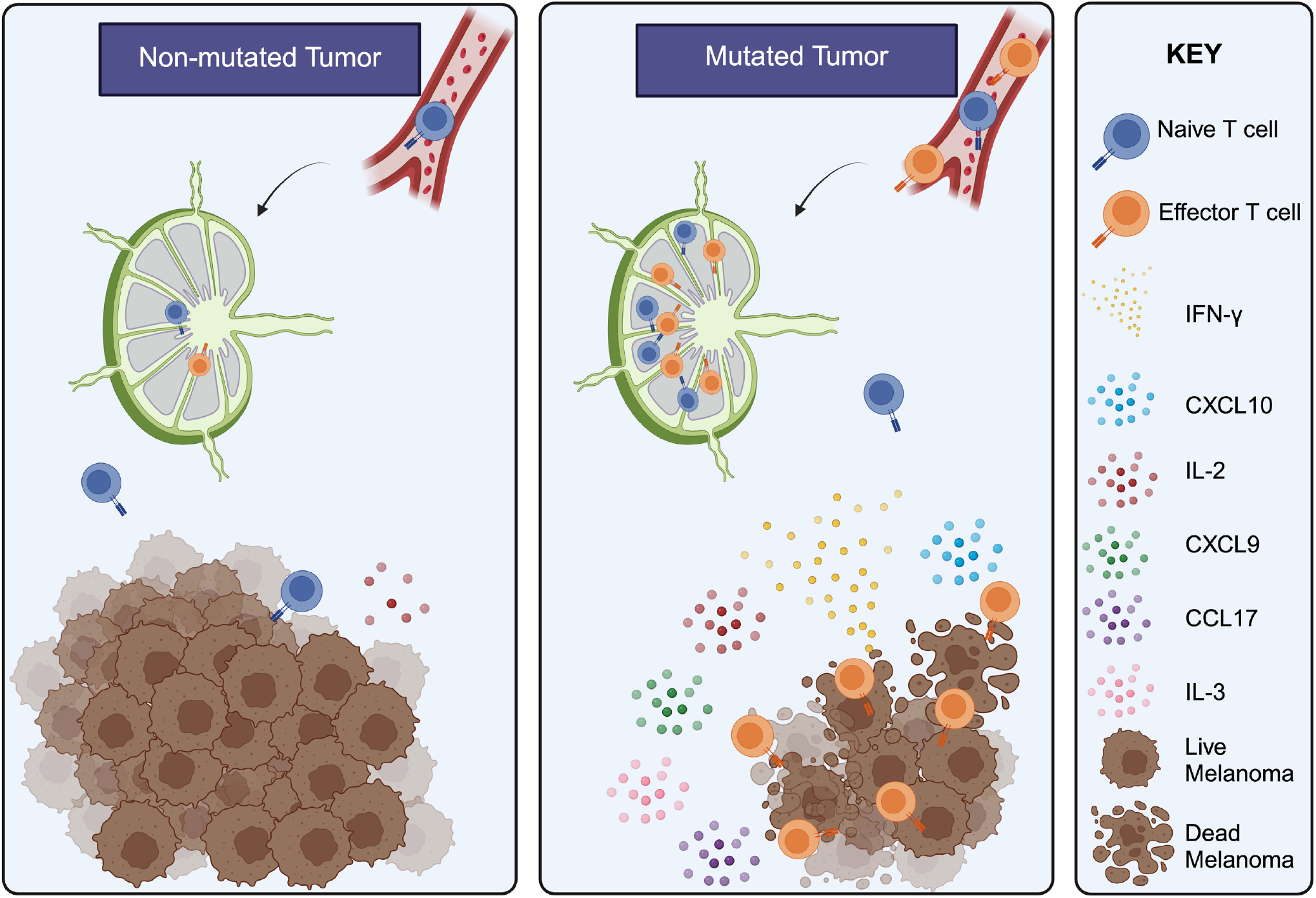
Summarized Results. **(A)** Visual depiction of main findings of the paper including increased cytokine and chemokine production, antitumor responses, and naïve to effector transition of adoptively transferred cells found in neoantigen tumors. Created using BioRender.

## Discussion

Using an antigen-specific adoptive cellular therapy model in melanoma, we uncovered striking differences in how an identical T cell product responds to neoantigen-expressing versus wild-type tumors. Adoptive transfer of naïve pmel-1 CD8 T cells elicited more potent antitumor responses against neoantigen-containing tumors, culminating in enhanced survival and protection from tumor recurrence. Genetic profiling of tumors revealed an immune stimulatory signature in neoantigen tumors that correlates with improved clinical outcomes. Moreover, distinct cytokine signatures emerged in response to neoantigen versus wild-type tumors, highlighting the nuanced nature of immune activation in these contexts.

Importantly, using naïve T cells in this model allowed us to systematically evaluate fundamental aspects of T cell activation and trafficking in the neoantigen versus wild-type setting. As exogenous IL-2 and vaccination were not needed for efficacy of this ACT product, we were able to delve into the intricate biology of the transferred cells in a very controlled setting. A key finding of our study centers on the differential trafficking and activation patterns of T cells across tumor types. While initial engraftment kinetics appeared similar between neoantigen and self-antigen groups, a significant divergence emerged over time. T cells responding to neoantigens demonstrated enhanced persistence in the tumor and across secondary lymphoid tissues. We propose that increased T cell avidity toward neoantigens, coupled with their escape from tolerance mechanisms that typically constrain responses to wild-type antigens, underlies this enhanced functionality(26,37-39).

Importantly, our findings challenge the traditional paradigm of T cell activation in cancer immunotherapy. Rather than simple tumor infiltration followed by local effector differentiation, we discovered a more complex progression. In mice bearing neoantigen-expressing tumors, naïve pmel-1 T cells underwent efficient lymph node homing followed by rapid effector differentiation, marked by CD44 upregulation and CD62L and TCF1 downregulation—a transition notably absent in wild-type tumor settings. Based on these observations, we propose a three-phase model of T cell activation in adoptive cell therapy: 1) Initial Lymphoid Homing (one to two days post-transfer): antigen-independent migration to secondary lymphoid organs, 2) Neoantigen-Driven Programming (three to four days post-ACT): early effector differentiation within lymph nodes and 3) Sustained Response (by ten days post-ACT): durable effector persistence across lymphoid tissues and tumors.

This early effector transition appears critical for establishing durable antitumor immunity, representing a distinctive feature of neoantigen recognition. The lymph node-derived differentiation and educational signals fundamentally shape T cell persistence within the tumor microenvironment. These insights carry important clinical implications, suggesting that strategies to enhance lymph node trafficking and neoantigen presentation—such as coordinated vaccine approaches—could augment ACT efficacy(18,40,41). Furthermore, our findings raise important considerations regarding the impact of surgical lymph node dissection on immunotherapy outcomes, warranting careful investigation in both preclinical and clinical settings.

Recent advances in genomic and proteomic technologies have accelerated neoantigen discovery(42-44), making our mechanistic insights particularly timely for advancing personalized cancer immunotherapy. The identification of shared, or “public,” neoantigens could enable broader therapeutic applications. Understanding the complex dynamics of T cell activation and differentiation in response to neoantigens remains crucial for optimizing immunotherapeutic strategies and improving patient outcomes in solid tumors. In conclusion, our study demonstrates that neoantigen recognition profoundly influences the activation, expansion, and persistence of adoptively transferred T cells, driving durable antitumor immunity. Understanding and leveraging these mechanisms will be critical for advancing neoantigen-targeted therapies and improving outcomes for patients with solid tumors.

## Acknowledgments

Research reported in this publication was supported in part by the Emory Integrated Genomics Core (EIGC), Winship Cancer Animal Models (CAMS) Core, and the Pediatrics/Winship Flow Cytometry Core Shared Resources of Winship Cancer Institute of Emory University and NIH/NCI under award number P30CA138292. The content is solely the responsibility of the authors and does not necessarily represent the official views of the National Institutes of Health.

## Author contributions

Conceptualization: H.M.K., C.M.P, and A.R.R.

Methodology: M.C.W., A.R.R, G.R.R., A.S.S., F.J.B., H.M.K

Investigation: M.C.W., A.R.R., M.M.W., A.C.C., M.B.W., A.T.R., S.K., H.M.K.

Visualization: M.C.W., A.R.R., A.C.C., H.M.K.

Supervision: C.M.P and G.B.L.

Writing—original draft: M.C.W., C.M.P., H.M.K.

Writing—review & editing: all authors

