## Supplemental Figures 1-5 and Supplemental Table 2 for "Neoantigens drive adoptively transferred CD8 T cells to long-lived effectors mediated by lymph node trafficking"

**Fig. S1.**

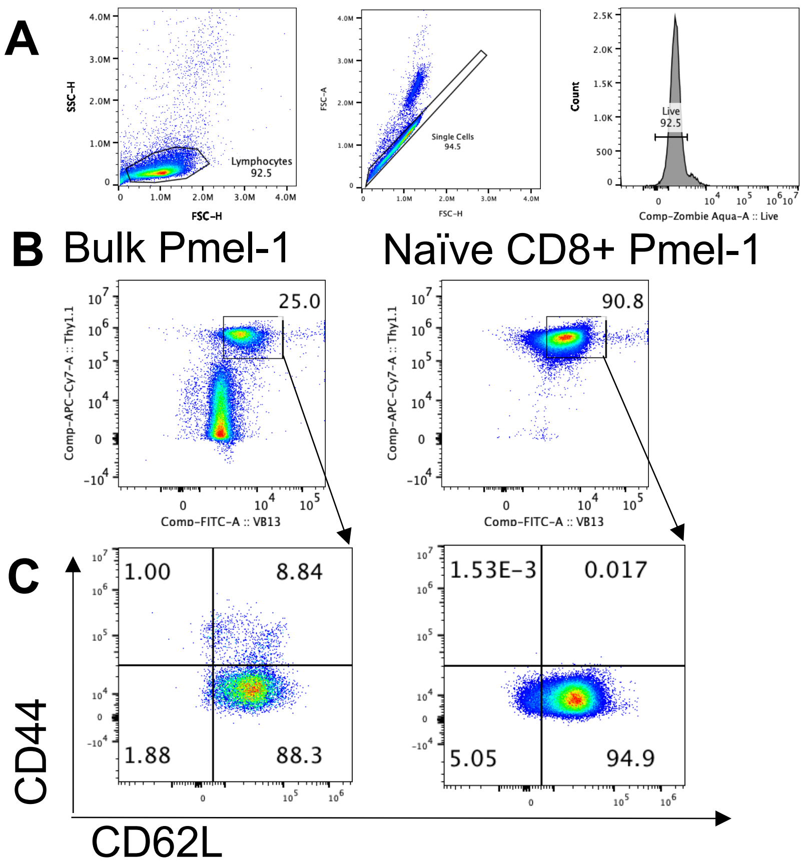


Supplemental Figure 1: Quality assurance by flow cytometry of naïve cytotoxic T cells prior adoptively transfer to mice bearing melanoma. (A) Gating Strategy of isolating lymphocytes followed by selection of singlets and subsequently live cells (B) Double positive Vb13+Thy1.1+ cells were gated on live cells (C) CD44 and CD62L characterization of Vb13+Thy1.1+ was used to assess efficacy of naïve T cell isolation(>94% yield).

### Fig. S2.

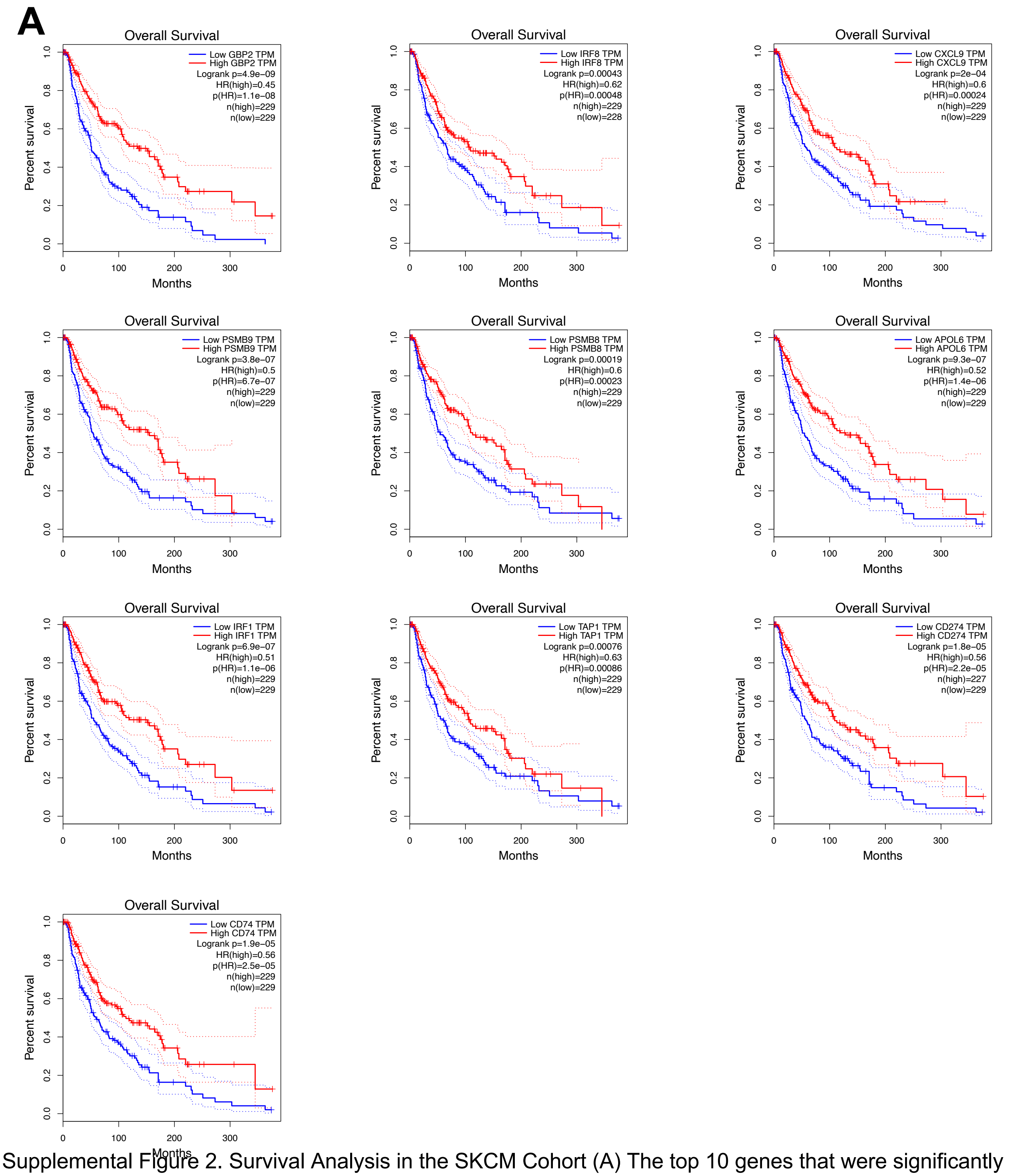


Supplemental Figure 2. Survival Analysis in the SKCM Cohort (A) The top 10 genes that were significantly upregulated in neoantigen tumors day 5 post-ACT were evaluated for their impact on survival in the TCGA SKCM cohort. Overall Survival with a Median Group Cutoff was used. Hazards Ratio was calculated based on the PH model. 95% confidence interval is displayed as a dotted line. Graphs made using GEPIA.

**Fig. S3.**

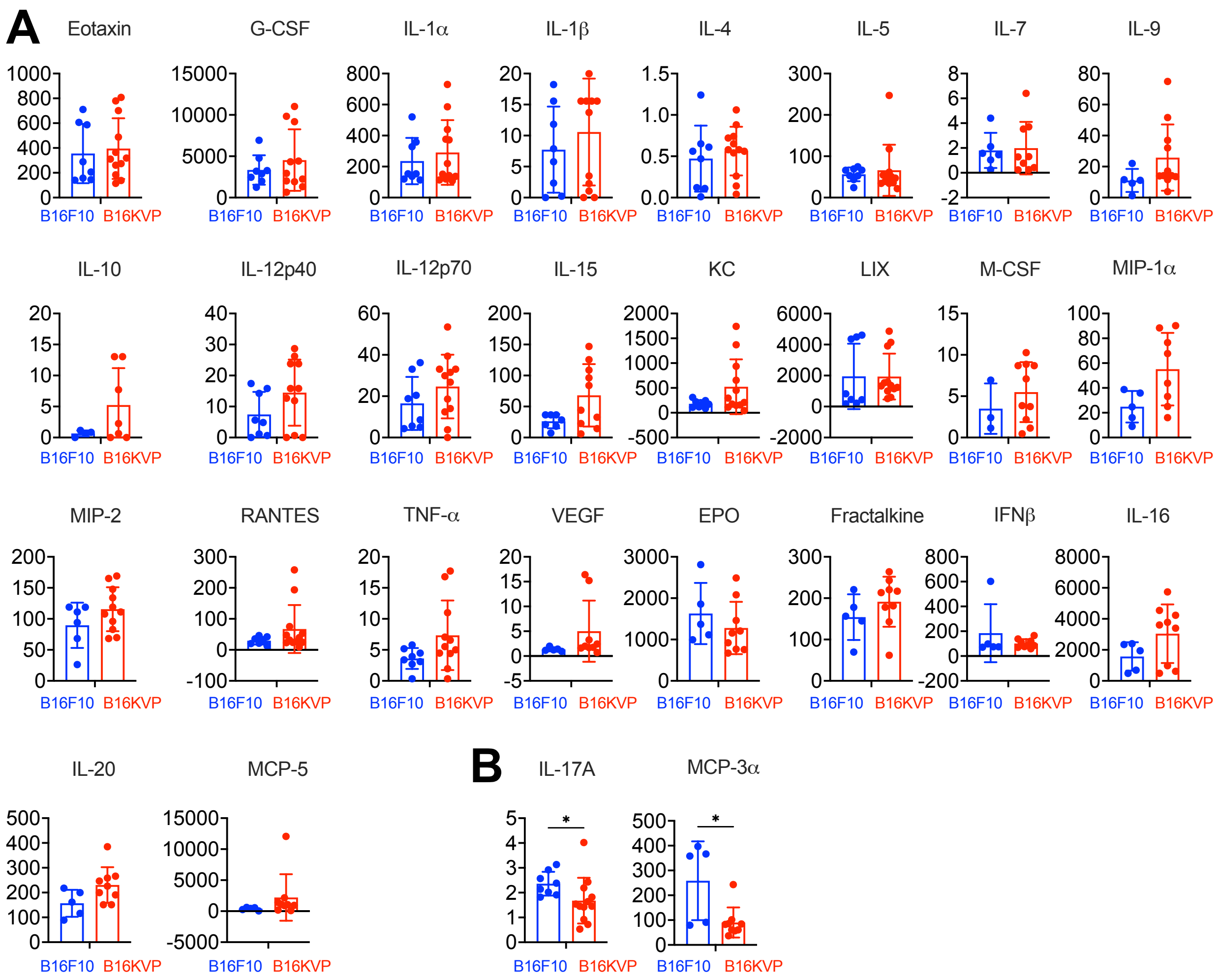


Supplemental Figure 3. Additional Cytokines (A) Cytokines that were not significantly different in serum of B16KVP versus B16F10 mice five days post-ACT. Mann-Whitney U test performed. (B) Cytokines significantly enriched in serum of B16F10 mice five days post-ACT. Mann-Whitney U test performed.

**Fig S4.**

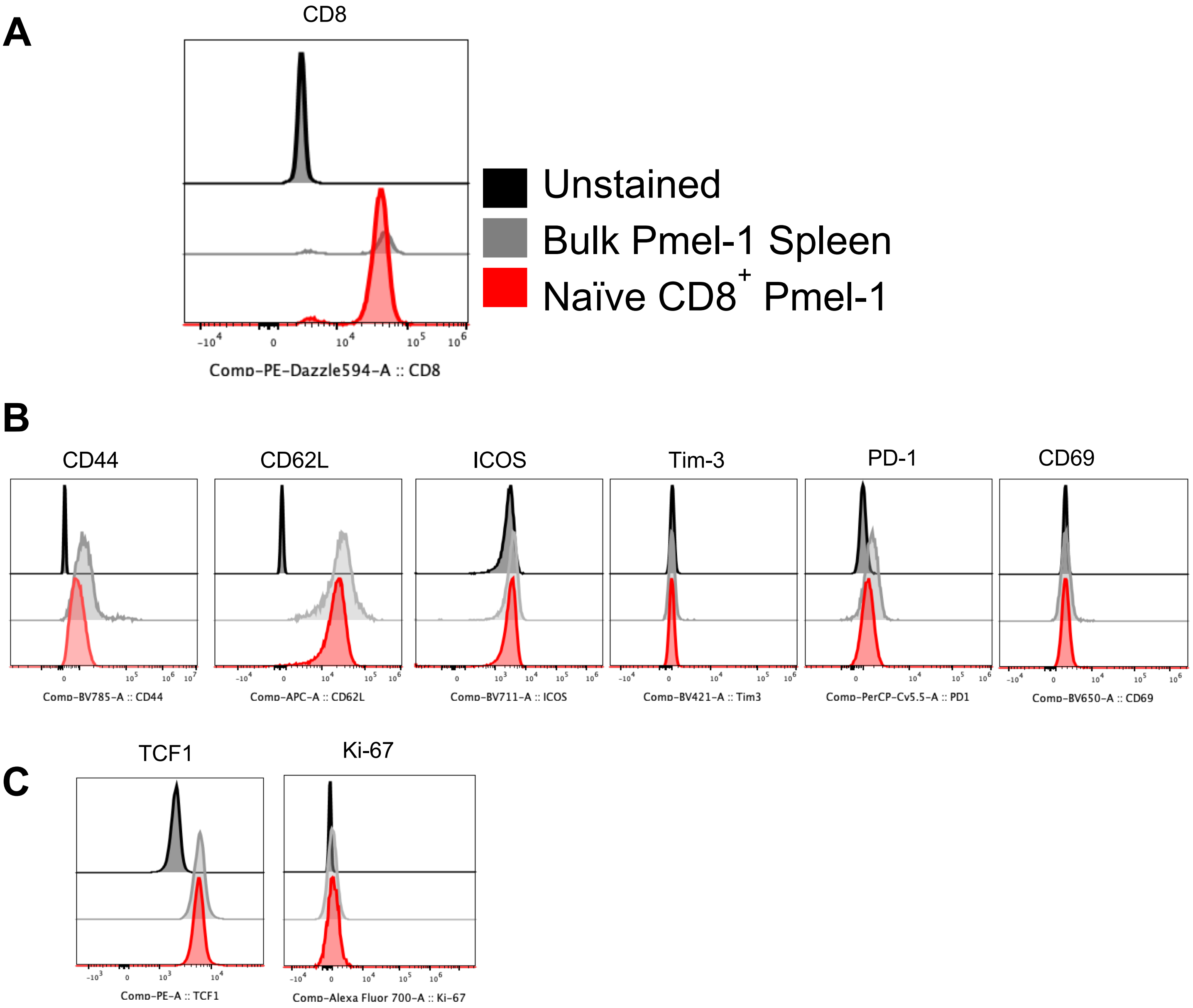


Supplemental Figure 4: Assessment of pmel-1 T cell phenotype prior to adoptive transfer (A) CD8 expression histogram on unstained, bulk pmel-1 cells or sorted naïve pmel-1 T cells. Gated on lymphocytes/single cell/live/CD45+ (B) Expression of surface markers CD44, CD62-L, ICOS, Tim-3, PD-1, and CD69 on unstained, bulk pmel-1 T cells, or sorted naïve pmel-1 T cells prior to adoptive transfer. Gated on Lymphocytes/Single Cells/Live/CD45+/Vβ13+ Thy1.1+. (C) Expression of intracellular markers TCF-1 and Ki-67 expression on unstained, bulk pmel-1 T cells, or naïve pmel-1 T cells prior to adoptive transfer.

**Fig S5.**

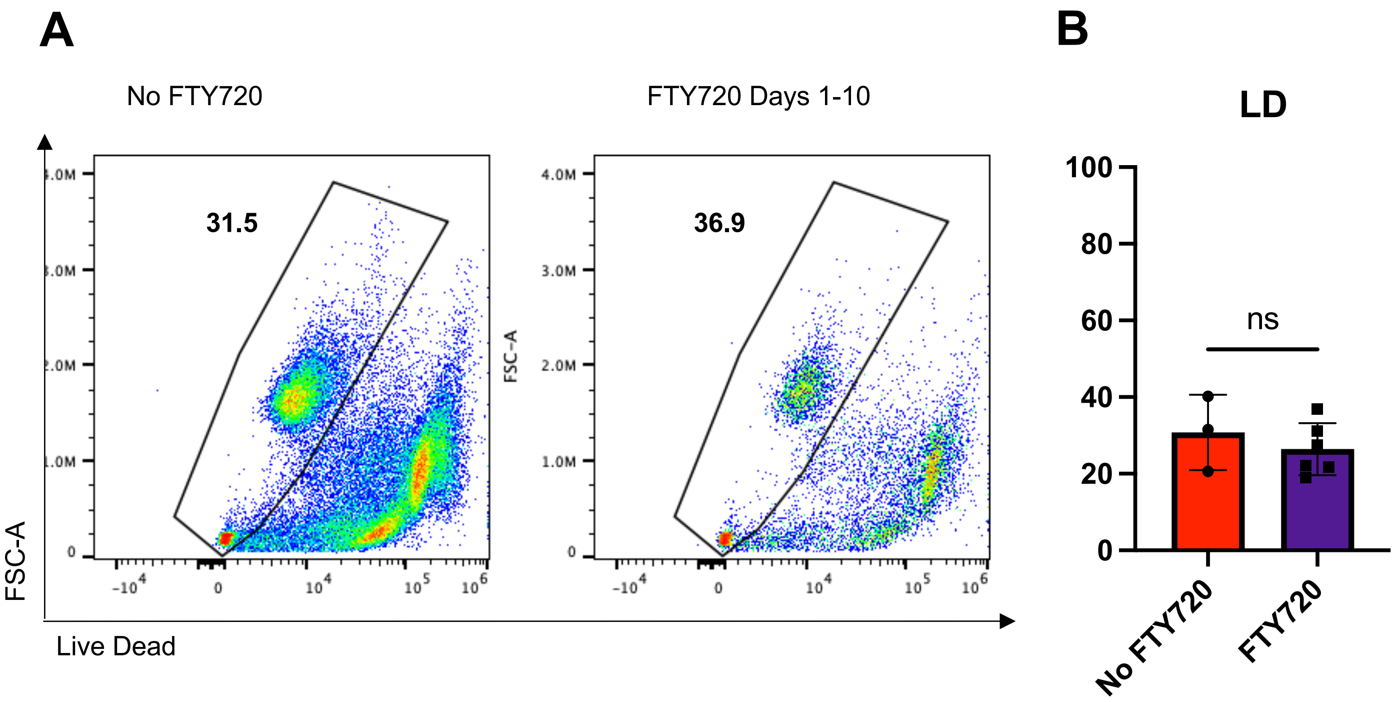


Supplemental Figure 5. FTY720 and Viability (A) Representative flow plots showing viability of cells in the lymph node of mice given FTY720 or no treatment. Gated on Lymphocytes/Single Cells. (B) Bar graph of viability for all samples. Mann-Whitney U test performed ($p=0.5476$).

**Table S1. (separate file)**

Table containing all statistically significant ($p < 0.05$) genes detected 5 days post-ACT from tumor tissue comparing mice with B16KVP to B16F10 tumors that received pmel-1 ACT.

**Table S2.**

Table of reagent information and antibodies used.

| Reagent or Resource | Source | Identifier |
|---|---|---|
| Antibodies: | | |
| Vβ13 | BD Biosciences | MR12-3 |
| Thy1.1 | Biolegend | Ox-7 |
| CD8a | Biolegend | 53-6.7 |
| CD44 | Biolegend | IM7 |
| CD62L | Biolegend | MEL-14 |
| CD69 | Biolegend | H1.2F3 |
| ICOS | Biolegend | C398.4A |
| Tim-3 | Biolegend | RMT3-23 |
| PD-1 | Biolegend | 29F.1A12 |
| T-bet | Biolegend | 4B10 |
| TCF1 | BD Biosciences | S33-966 |
| Ki-67 | Biolegend | 16A8 |
| Fixable Live Dead | ThermoFisher | L34957 |
| CD3e | BD Biosciences | 145-2C11 |
| CD45 | BD Biosciences | 30-F11 |
| Chemicals, Peptides, and Recombinant Proteins: | | |
| GP100 Peptide | Genscript | RP20344 |
| rhIL-2 | NIH Repository | |
| Fingolimod (FTY720) | Cayman Chemicals | 10006292 |
| RNAlater Solution | Invitrogen | AM7021 |
| 2 Mercaptoethanol 1000x | Fisher | 21985023 |
| 100mM Sodium Pyruvate Solution | Fisher | 11360-070 |
| HEPES | Fisher | 15630-080 |
| MEM non-essential amino acids | Fisher | SH30238.01 |
| Penicillin-Streptomycin (10,000 U/mL) | VWR | 45000-652 |
| Additional Reagants: | | |
| Microvette CB 300 K2E Tubes | SARSTEDT | 16.444.100 |
| Corning™ cellgro™ RPMI 1640 Medium (Mod.) 1X with L-Glutamine | VWR | MT10040CM |
| Critical Commercial Assays: | | |
| eBiosciences Foxp3/Transcription Factor Staining Kit | Thermo Fisher | 00-5523-00 |
| EasySep Mouse Naïve CD8+ T Cell Isolation Kit | Stemcell Technologies | 19858 |
| RBC Lysis Buffer | Biolegend | 420302 |
