## Supplemental Table 1 for "Neoantigens drive adoptively transferred CD8 T cells to long-lived effectors mediated by lymph node trafficking"

| Probe Name | B16KVP | StDev of KVP | B16F10 | StDev of B16F10 | B16KVP vs. B16F10 | P value of: B16KVP vs. B16F10 | t-statistic of: KVP vs. Reference | Lower 95% CI of: KVP vs. Reference | Upper 95% CI of: KVP vs. Reference | FDR adjusted pvalue of: KVP vs. Reference |
|---|---|---|---|---|---|---|---|---|---|---|
| Birc3 | 138.39 | 7.03 | 64.8 | 3.65 | 1.09 | 0.0000722 | 17.16286659 | 1.86 | 1.3 | 0.4 |
| Il6 | 62.96 | 6.05 | 19.1 | 1.31 | 1.72 | 0.00014539 | 17.30480576 | 2.65 | 2.04 | 0.4 |
| Cmtm6 | 620.13 | 44.33 | 277.93 | 9.36 | 1.16 | 0.0005094 | 17.75717735 | 1.84 | 1.44 | 0.52 |
| Irf9 | 660.8 | 39.78 | 207.27 | 25.16 | 1.67 | 0.0006838 | 14.98532391 | 2.29 | 2.15 | 0.52 |
| Axl | 579.46 | 87.2 | 117.2 | 26.26 | 2.31 | 0.00070014 | 10.70072556 | 3.07 | 2.99 | 0.52 |
| Lgals9 | 2106.21 | 211.89 | 776.29 | 109.01 | 1.44 | 0.00080898 | 10.13948154 | 1.98 | 1.89 | 0.52 |
| Ddb2 | 59.91 | 2.81 | 43.1 | 1.69 | 0.48 | 0.00081074 | 9.40380669 | 1.24 | 0.64 | 0.52 |
| Irf4 | 54.22 | 6.69 | 19.95 | 2.93 | 1.44 | 0.00084491 | 9.11864948 | 1.92 | 1.95 | 0.52 |
| Hdac4 | 219.54 | 9.88 | 161.06 | 6.34 | 0.45 | 0.00094998 | 8.9213829 | 1.22 | 0.61 | 0.52 |
| Cxcl10 | 3298.33 | 827.43 | 540.43 | 105.41 | 2.61 | 0.00097884 | 9.74557972 | 3.38 | 3.46 | 0.52 |
| Gbp3 | 655.49 | 209.16 | 66.15 | 20.26 | 3.31 | 0.00102112 | 8.6687746 | 4.27 | 4.52 | 0.52 |
| Ifit3 | 3055.26 | 482.34 | 409.97 | 129.13 | 2.9 | 0.00146514 | 10.59905529 | 4.08 | 3.77 | 0.66 |
| Rsad2 | 2191.35 | 655.68 | 396.48 | 77.91 | 2.47 | 0.00158063 | 8.45292759 | 2.9 | 3.4 | 0.66 |
| Kat2b | 729.87 | 57.88 | 407.77 | 42.12 | 0.84 | 0.00182975 | 7.75007915 | 1.41 | 1.18 | 0.66 |
| Myd88 | 349.71 | 57.62 | 129.73 | 11.7 | 1.43 | 0.00202332 | 9.30039215 | 1.92 | 1.92 | 0.66 |
| Oas1a | 1702.32 | 387.1 | 409.15 | 49.79 | 2.06 | 0.00206746 | 9.83517933 | 2.62 | 2.72 | 0.66 |
| Cd47 | 1309.61 | 249.83 | 447.23 | 82.49 | 1.55 | 0.00231104 | 7.01377916 | 1.8 | 2.25 | 0.66 |
| Isg15 | 2636.95 | 359.4 | 469.93 | 135.67 | 2.49 | 0.00231594 | 10.0602026 | 2.68 | 3.55 | 0.66 |
| Ly6c1 | 3889.51 | 1241.3 | 562.04 | 221.25 | 2.79 | 0.00234856 | 6.87697268 | 2.83 | 4.08 | 0.66 |
| Icam1 | 260.07 | 88.22 | 36.62 | 13.82 | 2.83 | 0.00241435 | 6.86280727 | 2.86 | 4.14 | 0.66 |
| Deptor | 722.23 | 77.41 | 187.46 | 41.24 | 1.95 | 0.00247951 | 9.878479 | 2.14 | 2.79 | 0.66 |
| Nfkbia | 813.94 | 130.85 | 309.1 | 22.44 | 1.4 | 0.00301947 | 9.69421959 | 1.71 | 2.02 | 0.69 |
| Mill2 | 159.21 | 32.23 | 40.52 | 3.73 | 1.97 | 0.00312766 | 10.25992012 | 2.21 | 2.8 | 0.69 |
| Ccne1 | 327.88 | 15.96 | 448.13 | 28.42 | -0.45 | 0.00321338 | -6.76685762 | 0.63 | -0.24 | 0.69 |
| Ccl2 | 3281.25 | 1000.9 | 693.91 | 196.78 | 2.24 | 0.00346717 | 6.46219301 | 2.2 | 3.35 | 0.69 |
| Msh6 | 639.56 | 28.27 | 836.12 | 47.37 | -0.39 | 0.00358821 | -6.49080181 | 0.67 | -0.2 | 0.69 |
| Esr1 | 141.97 | 38.22 | 30.98 | 9.48 | 2.2 | 0.00382793 | 6.09573412 | 2.07 | 3.34 | 0.69 |
| H2-M3 | 196.63 | 71.96 | 31.54 | 11.12 | 2.64 | 0.00384148 | 6.18366289 | 2.43 | 4 | 0.69 |
| Tnfaip3 | 147.52 | 20.89 | 70.4 | 10.98 | 1.07 | 0.00388144 | 6.10493469 | 1.42 | 1.62 | 0.69 |
| Uba7 | 422.67 | 116.11 | 88.96 | 31.3 | 2.25 | 0.00395821 | 6.05941057 | 2.1 | 3.43 | 0.69 |
| Adm | 307.49 | 35.92 | 652.85 | 29.09 | -1.09 | 0.003985 | -10.23190403 | 0.34 | -0.63 | 0.69 |
| Ifi203 | 638.2 | 132.13 | 85.5 | 35.38 | 2.9 | 0.00408069 | 7.87605381 | 3.31 | 4.07 | 0.69 |
| Ctss | 1810.34 | 561.42 | 355.71 | 123.7 | 2.35 | 0.00412489 | 5.9511528 | 2.13 | 3.6 | 0.69 |
| Ifit1 | 446.18 | 112.65 | 113.5 | 35.56 | 1.97 | 0.00422731 | 6.05866098 | 1.92 | 3.01 | 0.69 |
| Tlr3 | 318.8 | 63.33 | 123.22 | 17.51 | 1.37 | 0.0045128 | 6.58723831 | 1.63 | 2.03 | 0.72 |
| Parp4 | 331.77 | 32.21 | 160.35 | 24.62 | 1.05 | 0.00479334 | 6.74093294 | 1.47 | 1.54 | 0.74 |
| Cxcl1 | 106.5 | 18.72 | 23.66 | 8.42 | 2.17 | 0.00515793 | 7.17706394 | 2.31 | 3.13 | 0.77 |
| Parp9 | 1006.89 | 375.97 | 178.32 | 55.23 | 2.5 | 0.00522894 | 6.01203346 | 2.26 | 3.82 | 0.77 |
| B2m | 46116.75 | 15686.79 | 8619.28 | 1808.91 | 2.42 | 0.00561146 | 6.98003387 | 2.49 | 3.52 | 0.78 |
| Zc3h12a | 184.67 | 23.08 | 98.75 | 4.75 | 0.9 | 0.00573805 | 8.35012436 | 1.35 | 1.37 | 0.78 |
| Csf1 | 403.02 | 76.38 | 104.27 | 36.01 | 1.95 | 0.00589249 | 6.52927828 | 2 | 2.9 | 0.78 |
| Pfkfb3 | 470.46 | 76.78 | 248.57 | 32.38 | 0.92 | 0.00592502 | 5.51121902 | 1.31 | 1.45 | 0.78 |
| Oas3 | 102.49 | 18.49 | 25.5 | 8.75 | 2.01 | 0.00610959 | 6.45459747 | 2.02 | 3 | 0.79 |
| Fcgr4 | 783.85 | 344.5 | 107.33 | 33.38 | 2.87 | 0.00628233 | 6.18251085 | 2.62 | 4.35 | 0.79 |
| Cxcl9 | 2482.02 | 1792.67 | 112.5 | 68.58 | 4.46 | 0.00671162 | 5.23097229 | 3.36 | 7.18 | 0.79 |
| Stc1 | 392.89 | 83.58 | 130.52 | 37.23 | 1.59 | 0.00677837 | 5.50387669 | 1.59 | 2.51 | 0.79 |
| Lilrb4a | 753.1 | 198.19 | 244.2 | 74.13 | 1.62 | 0.00680722 | 5.15221405 | 1.54 | 2.63 | 0.79 |
| Syk | 166.65 | 29.15 | 72.49 | 14.52 | 1.2 | 0.00690524 | 5.18467617 | 1.38 | 1.94 | 0.79 |
| Fcgr3 | 147.97 | 43.88 | 35.46 | 13.81 | 2.06 | 0.00696603 | 5.20468807 | 1.74 | 3.32 | 0.79 |
| Cebpb | 667.19 | 71.91 | 244.98 | 53.04 | 1.45 | 0.00717792 | 6.83470869 | 1.45 | 2.36 | 0.8 |
| Pla1a | 128.63 | 39.75 | 30.8 | 12.65 | 2.06 | 0.00733075 | 5.05871582 | 1.7 | 3.36 | 0.8 |
| Stat2 | 915.22 | 331.5 | 120.32 | 11.11 | 2.93 | 0.00766403 | 9.25526524 | 2.96 | 4.29 | 0.82 |
| Sfrp1 | 381 | 57.2 | 63.68 | 28.09 | 2.58 | 0.00831594 | 7.40590811 | 2.12 | 4.08 | 0.83 |
| Oas2 | 332.95 | 169.36 | 62.26 | 16.31 | 2.42 | 0.00836482 | 5.66727877 | 2.09 | 3.78 | 0.83 |
| Tnfsf10 | 126.28 | 35.12 | 29.01 | 11.36 | 2.12 | 0.00839536 | 5.28861523 | 1.8 | 3.4 | 0.83 |
| Cd300a | 99.97 | 36.48 | 23.66 | 8.42 | 2.08 | 0.0087035 | 4.93273163 | 1.67 | 3.42 | 0.83 |
| Relb | 243.76 | 33.96 | 143.85 | 13.25 | 0.76 | 0.00876716 | 5.36256075 | 1.24 | 1.21 | 0.83 |
| Bcl2l1 | 808.08 | 24.5 | 524.5 | 48.69 | 0.62 | 0.00882656 | 7.7979126 | 1.21 | 0.97 | 0.83 |
| Hmgb1 | 2426.48 | 160.83 | 3390.91 | 320.47 | -0.48 | 0.00891035 | -5.13599062 | 0.58 | -0.18 | 0.83 |
| Ptpn11 | 829.65 | 27.38 | 536.67 | 51.98 | 0.63 | 0.00912537 | 7.50193071 | 1.2 | 0.99 | 0.83 |
| Ifit2 | 1727.01 | 718.42 | 344.97 | 80.44 | 2.32 | 0.00920266 | 5.65023708 | 2.02 | 3.63 | 0.83 |
| Nos2 | 168.53 | 51.97 | 51.08 | 14.44 | 1.72 | 0.00923802 | 4.71914387 | 1.48 | 2.88 | 0.83 |
| H2-K1 | 9435.27 | 4206.64 | 1523.45 | 408.56 | 2.63 | 0.00992868 | 5.67918682 | 2.23 | 4.1 | 0.85 |
| Ifih1 | 857.6 | 126.77 | 161.04 | 59.1 | 2.41 | 0.00998409 | 6.84793234 | 1.86 | 3.93 | 0.85 |
| Ifnar1 | 818.1 | 44.25 | 646.56 | 42.85 | 0.34 | 0.01021627 | 4.70794296 | 1.08 | 0.57 | 0.85 |
| Dnajc14 | 363.65 | 25.4 | 277.08 | 9.48 | 0.39 | 0.01024987 | 6.03213072 | 1.08 | 0.67 | 0.85 |
| Il15 | 56 | 16.31 | 21.89 | 4.72 | 1.36 | 0.01083862 | 4.70906258 | 1.36 | 2.27 | 0.85 |
| Ripk3 | 112.39 | 18.75 | 41.47 | 12.96 | 1.44 | 0.01089873 | 5.18546438 | 1.47 | 2.32 | 0.85 |
| Ikzf2 | 135.01 | 47.07 | 37.5 | 13.77 | 1.85 | 0.01103375 | 4.50998497 | 1.46 | 3.15 | 0.85 |
| Oaz1 | 3395.33 | 153.48 | 4593.16 | 391.42 | -0.44 | 0.01103846 | -5.52777863 | 0.62 | -0.18 | 0.85 |
| Mx1 | 423.93 | 19.18 | 105.46 | 31.66 | 2.01 | 0.01111928 | 8.66148663 | 2.01 | 3 | 0.85 |
| Gbp2 | 10070.27 | 10140.43 | 316.87 | 207.43 | 4.99 | 0.01124138 | 4.95710278 | 3.45 | 8.19 | 0.85 |
| Lif | 142.8 | 46.23 | 44.66 | 15.07 | 1.68 | 0.01129484 | 4.5032568 | 1.41 | 2.86 | 0.85 |
| Irf7 | 805.01 | 68.77 | 147.88 | 60.95 | 2.44 | 0.01155869 | 7.89789009 | 2.16 | 3.78 | 0.85 |
| Notch1 | 281.32 | 81.52 | 79.28 | 7.53 | 1.83 | 0.01182859 | 6.98789263 | 1.63 | 2.95 | 0.85 |
| Cdk6 | 329.15 | 75.56 | 147.49 | 31.72 | 1.16 | 0.01217627 | 4.3805356 | 1.25 | 2 | 0.85 |
| Psmb8 | 4661.11 | 3480.92 | 236.44 | 82.3 | 4.3 | 0.01226472 | 5.76660109 | 2.13 | 7.51 | 0.85 |
| H2-Ab1 | 2652.87 | 1535.93 | 284.46 | 67.22 | 3.22 | 0.01239095 | 6.13176489 | 1.95 | 5.48 | 0.85 |
| H2-Eb1 | 633.25 | 382.59 | 62.83 | 15.46 | 3.33 | 0.01239756 | 6.17712975 | 2.02 | 5.65 | 0.85 |
| Tap2 | 666.15 | 378.52 | 82.77 | 28.59 | 3.01 | 0.01250472 | 5.02665329 | 2.15 | 4.91 | 0.85 |
| H2-D1 | 9592.5 | 3797.67 | 1533.77 | 197.7 | 2.64 | 0.01252729 | 7.06978893 | 2.05 | 4.25 | 0.85 |
| Tnfrsf14 | 255.86 | 147.56 | 21.86 | 4.03 | 3.55 | 0.01255316 | 6.87072992 | 2.51 | 5.77 | 0.85 |
| Ccl7 | 1482.48 | 622.55 | 334.77 | 128.54 | 2.15 | 0.01276756 | 4.52167559 | 1.55 | 3.66 | 0.86 |
| Jak2 | 271.37 | 37.46 | 149.56 | 29.3 | 0.86 | 0.01317961 | 4.42658091 | 1.18 | 1.48 | 0.86 |
| Lamc2 | 52.16 | 9.42 | 27.38 | 5.56 | 0.93 | 0.0136152 | 4.21073818 | 1.17 | 1.63 | 0.86 |
| Fas | 90.94 | 13.55 | 28.06 | 9.71 | 1.7 | 0.01369787 | 5.67269373 | 1.33 | 2.98 | 0.86 |
| Il33 | 377.31 | 143.76 | 102.48 | 17.4 | 1.88 | 0.01373422 | 5.53599548 | 1.34 | 3.34 | 0.86 |
| Parp12 | 2589.2 | 1042.48 | 479.14 | 56.73 | 2.43 | 0.01388228 | 6.77400875 | 1.85 | 3.98 | 0.86 |
| Thbs1 | 575.59 | 189.11 | 205.29 | 57.96 | 1.49 | 0.01406549 | 4.27037001 | 1.3 | 2.6 | 0.86 |
| Cd40 | 58.74 | 21.22 | 20.75 | 4.61 | 1.5 | 0.01437141 | 4.63812733 | 1.39 | 2.53 | 0.86 |
| Itgb2 | 281.74 | 116.28 | 77.31 | 26.68 | 1.87 | 0.01456927 | 4.28550339 | 1.39 | 3.25 | 0.86 |
| Snca | 1630.86 | 158.52 | 1128.98 | 128.05 | 0.53 | 0.01461222 | 4.22893429 | 1.1 | 0.93 | 0.86 |
| Il18 | 318.83 | 69.86 | 155.9 | 20.53 | 1.03 | 0.01473252 | 4.74408054 | 1.27 | 1.72 | 0.86 |
| Socs1 | 566.23 | 358.76 | 48.02 | 12.56 | 3.56 | 0.0151413 | 5.80897665 | 1.9 | 6.2 | 0.86 |
| Egr1 | 282.19 | 71.95 | 120.76 | 28.62 | 1.22 | 0.01520597 | 4.07557297 | 1.2 | 2.18 | 0.86 |
| Oasl1 | 170.08 | 30.81 | 70.85 | 19.47 | 1.26 | 0.01540826 | 4.45337534 | 1.28 | 2.17 | 0.86 |
| H2-T23 | 3104.59 | 2040.84 | 274.2 | 100.5 | 3.5 | 0.01545983 | 5.23985815 | 1.54 | 6.38 | 0.86 |
| Vcam1 | 118.92 | 27.68 | 58.05 | 11.86 | 1.03 | 0.01568858 | 4.09606504 | 1.17 | 1.84 | 0.86 |
| Nbn | 545.47 | 6.12 | 685.39 | 37.39 | -0.33 | 0.01576493 | -7.03822517 | 0.69 | -0.13 | 0.86 |
| Cd74 | 4097.54 | 2635.76 | 335.41 | 68.31 | 3.61 | 0.01583184 | 6.17236853 | 2.13 | 6.13 | 0.86 |
| Stat1 | 296.59 | 180.54 | 40.22 | 14.6 | 2.88 | 0.01588272 | 4.60856199 | 1.86 | 4.87 | 0.86 |
| Ifi35 | 1965.55 | 942.98 | 372.42 | 80.52 | 2.4 | 0.01593795 | 5.3580637 | 1.39 | 4.33 | 0.86 |
| Angpt1 | 96.97 | 26.93 | 38.27 | 11.2 | 1.34 | 0.0160164 | 4.0344758 | 1.22 | 2.4 | 0.86 |
| Nlrc5 | 1119.67 | 579.73 | 95.03 | 5.08 | 3.56 | 0.0161342 | 7.57643318 | 2.9 | 5.58 | 0.86 |
| Slc11a1 | 314.85 | 152.9 | 79.51 | 17.68 | 1.99 | 0.01629607 | 4.92649984 | 1.19 | 3.72 | 0.86 |
| Tapbp | 946.93 | 495.58 | 130.6 | 28.24 | 2.86 | 0.01665992 | 5.58634233 | 1.58 | 5.06 | 0.87 |
| Irf5 | 243.04 | 114.4 | 58.46 | 24.01 | 2.06 | 0.01682633 | 4.01199198 | 1.34 | 3.69 | 0.87 |
| Ccnd1 | 2427.26 | 274.56 | 1678.72 | 202.52 | 0.53 | 0.01686256 | 3.95001459 | 1.07 | 0.96 | 0.87 |
| Cd84 | 215.9 | 68.12 | 71.92 | 27.56 | 1.59 | 0.01711388 | 3.93011022 | 1.23 | 2.87 | 0.87 |
| Casp1 | 273.42 | 113.49 | 78.57 | 12.75 | 1.8 | 0.01737746 | 5.22848082 | 1.25 | 3.28 | 0.88 |
| Cxcl16 | 223.08 | 65.05 | 72.42 | 6.37 | 1.62 | 0.01808763 | 6.06911325 | 1.39 | 2.77 | 0.89 |
| H2afx | 875.18 | 42.05 | 1048.17 | 65.7 | -0.26 | 0.01809251 | -3.98679543 | 0.72 | -0.05 | 0.89 |
| Psmb9 | 3572.89 | 3131.58 | 169.63 | 64.23 | 4.4 | 0.01827882 | 5.25220108 | 1.73 | 8 | 0.89 |
| Tnfrsf1a | 1696.2 | 336.69 | 900.96 | 171.83 | 0.91 | 0.01829056 | 3.86275172 | 1.12 | 1.66 | 0.89 |
| Il2rg | 283.72 | 133.03 | 87.88 | 27.65 | 1.69 | 0.01933588 | 4.05873775 | 1.29 | 3.02 | 0.93 |
| Nfil3 | 325.03 | 74.66 | 174.65 | 33.01 | 0.9 | 0.01943994 | 3.84608698 | 1.11 | 1.64 | 0.93 |
| Tap1 | 2903.21 | 1993.43 | 196.81 | 38.64 | 3.88 | 0.01949166 | 6.05225372 | 2.18 | 6.64 | 0.93 |
| Itgam | 271.18 | 101.43 | 86.69 | 21.77 | 1.65 | 0.02011515 | 4.20909882 | 1.32 | 2.89 | 0.95 |
| Dtx3l | 1264.01 | 614.35 | 228.92 | 37.23 | 2.47 | 0.02063064 | 5.44472504 | 1.43 | 4.41 | 0.96 |
| Tnfsf12 | 78.35 | 16.66 | 42.05 | 8.95 | 0.9 | 0.02081021 | 3.70441937 | 1.09 | 1.67 | 0.96 |
| Ccr5 | 158.56 | 69.98 | 39.39 | 6.53 | 2.01 | 0.02107152 | 5.06970501 | 1.23 | 3.71 | 0.96 |
| Kit | 118.11 | 26.42 | 60.75 | 14.99 | 0.96 | 0.0213165 | 3.68573666 | 1.1 | 1.79 | 0.96 |
| Ripk1 | 314.25 | 66.43 | 171.47 | 29.18 | 0.87 | 0.02169215 | 3.83766913 | 1.11 | 1.6 | 0.96 |
| Tnfrsf1b | 71.52 | 20.83 | 30.94 | 4.41 | 1.21 | 0.02212517 | 4.51636028 | 1.04 | 2.36 | 0.96 |
| 4930578C19Rik | 23.22 | 1.46 | 19.1 | 1.31 | 0.28 | 0.02212541 | 3.64132309 | 1.02 | 0.53 | 0.96 |
| Snai1 | 23.22 | 1.46 | 19.1 | 1.31 | 0.28 | 0.02212541 | 3.64132309 | 1.02 | 0.53 | 0.96 |
| Irf1 | 4533.29 | 3833.74 | 270.84 | 96.93 | 4.07 | 0.02224538 | 5.07972765 | 1.54 | 7.51 | 0.96 |
| Batf3 | 33.09 | 6.23 | 19.1 | 1.31 | 0.79 | 0.02224802 | 4.98415852 | 1.08 | 1.48 | 0.96 |
| Trim21 | 503.37 | 197.14 | 116.22 | 11.08 | 2.11 | 0.022297 | 5.84667969 | 1.47 | 3.67 | 0.96 |
| Tfrc | 1595.91 | 63.59 | 3695.24 | 939.68 | -1.21 | 0.02260695 | -6.08794737 | 0.24 | -0.36 | 0.96 |
| Melk | 260.22 | 36.83 | 386.92 | 36.92 | -0.57 | 0.02281171 | -3.9657259 | 0.49 | -0.11 | 0.96 |
| Cxcl5 | 161.42 | 3.24 | 29.9 | 14.19 | 2.43 | 0.02291997 | 6.45821619 | 1.76 | 4.05 | 0.96 |
| Ikbkb | 249.29 | 15.79 | 174.53 | 21.55 | 0.51 | 0.02301511 | 4.39829969 | 1.01 | 1.02 | 0.96 |
| Clec4e | 118.86 | 42.77 | 47.44 | 9.12 | 1.33 | 0.02359594 | 4.0533843 | 1.22 | 2.37 | 0.97 |
| Tapbpl | 200.52 | 132.96 | 26.59 | 9.56 | 2.91 | 0.02391863 | 4.38895845 | 1.04 | 5.77 | 0.97 |
| Cd274 | 1936.73 | 1379.49 | 143.33 | 25.56 | 3.76 | 0.02432514 | 5.59802198 | 1.83 | 6.64 | 0.97 |
| Rbl2 | 406.33 | 15.93 | 315.7 | 28.23 | 0.36 | 0.02441354 | 4.52172613 | 1.01 | 0.71 | 0.97 |
| Tgfb1 | 581.43 | 86.9 | 379.62 | 57.25 | 0.62 | 0.02445137 | 3.52023911 | 1.04 | 1.17 | 0.97 |
| Cd68 | 747.14 | 207.57 | 334.59 | 93.29 | 1.16 | 0.02449997 | 3.5543313 | 1.09 | 2.2 | 0.97 |
| Csf2rb | 94.89 | 23.89 | 34.07 | 13.2 | 1.48 | 0.02493796 | 3.84917569 | 1.19 | 2.7 | 0.97 |
| Csf3r | 57.18 | 20.93 | 19.1 | 1.31 | 1.58 | 0.02521358 | 5.61228752 | 1.29 | 2.8 | 0.97 |
| Itgb3 | 154.79 | 13.51 | 119.71 | 10.76 | 0.37 | 0.0252167 | 3.48882771 | 1.02 | 0.71 | 0.97 |
| Ptgs2 | 282.63 | 210.49 | 60.05 | 24.17 | 2.23 | 0.0252228 | 3.87005734 | 1.32 | 4.07 | 0.97 |
| Hdc | 44.93 | 11.31 | 19.1 | 1.31 | 1.23 | 0.02529925 | 5.36172819 | 1.18 | 2.22 | 0.97 |
| Cybb | 250.91 | 135.62 | 54.58 | 31.6 | 2.2 | 0.0253944 | 3.47972226 | 1.14 | 4.21 | 0.97 |
| Psmb10 | 1157.93 | 824.95 | 136.31 | 46.97 | 3.09 | 0.02540783 | 4.29559708 | 1 | 6.18 | 0.97 |
| Entpd1 | 93.11 | 34.65 | 29.42 | 12.26 | 1.66 | 0.02578313 | 3.46392179 | 1.1 | 3.19 | 0.98 |
| Maml2 | 231.23 | 65.76 | 95.51 | 31.49 | 1.28 | 0.02614937 | 3.45189357 | 1.07 | 2.45 | 0.98 |
| Vsir | 148.57 | 67.73 | 39.42 | 19.93 | 1.91 | 0.02621807 | 3.4746964 | 1.12 | 3.67 | 0.98 |
| Ngfr | 23.92 | 2.05 | 19.1 | 1.31 | 0.32 | 0.0266337 | 3.52998137 | 1.02 | 0.62 | 0.98 |
| Rb1 | 314.23 | 40.31 | 156.05 | 40.74 | 1.01 | 0.02673484 | 4.12250566 | 0.97 | 2.06 | 0.98 |
| Cd1d1 | 90.99 | 49.16 | 19.1 | 1.31 | 2.25 | 0.02712072 | 5.66732359 | 1.46 | 3.96 | 0.99 |
| Itga6 | 467.37 | 109.72 | 265.87 | 46.07 | 0.81 | 0.02763909 | 3.53036571 | 1.06 | 1.55 | 1 |
| Lair1 | 64.46 | 27.22 | 23.24 | 7.37 | 1.47 | 0.02834104 | 3.54454637 | 1.11 | 2.79 | 1 |
| H2-Aa | 3341.05 | 2602.3 | 318.77 | 46.15 | 3.39 | 0.0295121 | 5.25497484 | 1.53 | 6.17 | 1 |
| Sell | 93.3 | 56.49 | 19.1 | 1.31 | 2.29 | 0.02966462 | 5.44310617 | 1.39 | 4.1 | 1 |
| Ccl3 | 123.63 | 50.1 | 41.5 | 15.75 | 1.57 | 0.029685 | 3.40943623 | 1.08 | 3.05 | 1 |
| Nfkbie | 117.63 | 33.25 | 57.79 | 7.9 | 1.03 | 0.03029681 | 3.97799325 | 0.94 | 2.13 | 1 |
| Ccl19 | 201.68 | 88.52 | 64.43 | 20.81 | 1.65 | 0.03063935 | 3.50512028 | 1.11 | 3.14 | 1 |
| Spp1 | 2274.67 | 533.79 | 971.59 | 382.74 | 1.23 | 0.03071004 | 3.55686712 | 1.09 | 2.33 | 1 |
| Akt1 | 2071.17 | 76.93 | 3116.32 | 430.35 | -0.59 | 0.03149383 | -4.81541157 | 0.46 | -0.06 | 1 |
| H2-DMb1 | 230.8 | 175.69 | 27.7 | 9.67 | 3.06 | 0.03150973 | 4.02513504 | 0.86 | 6.33 | 1 |
| Irf2 | 363.72 | 92.26 | 187.82 | 27.88 | 0.95 | 0.03155952 | 3.71328497 | 1.1 | 1.77 | 1 |
| Fcgr2b | 1087.05 | 168.68 | 357.71 | 165.14 | 1.6 | 0.03157367 | 4.28105688 | 0.99 | 3.22 | 1 |
| Olfml2b | 233.64 | 60.68 | 96.62 | 40.3 | 1.27 | 0.03174271 | 3.45234179 | 1.07 | 2.45 | 1 |
| Exo1 | 121.26 | 18.32 | 183.8 | 8.5 | -0.6 | 0.03188121 | -4.57672787 | 0.45 | -0.04 | 1 |
| Tlr2 | 68.73 | 12.3 | 24.85 | 11.59 | 1.47 | 0.0321918 | 3.9616828 | 0.92 | 3.06 | 1 |
| Pirb | 138.09 | 71.47 | 35.53 | 19.95 | 1.96 | 0.03233006 | 3.25285435 | 1.03 | 3.87 | 1 |
| Polr2a | 1365.81 | 63.93 | 1151.56 | 81.76 | 0.25 | 0.03324764 | 3.44538116 | 1.01 | 0.47 | 1 |
| Hck | 61.82 | 25.65 | 19.1 | 1.31 | 1.69 | 0.03376633 | 4.99233723 | 1.18 | 3.16 | 1 |
| Gimap6 | 87.4 | 17.74 | 52.11 | 5.39 | 0.75 | 0.03471743 | 3.76880145 | 0.93 | 1.6 | 1 |
| Sbno2 | 188.01 | 24.85 | 127.11 | 21.95 | 0.56 | 0.03515207 | 3.22591901 | 1.01 | 1.12 | 1 |
| Cdh11 | 22.72 | 1.6 | 19.1 | 1.31 | 0.25 | 0.03587802 | 3.11010504 | 1 | 0.51 | 1 |
| Ifi27 | 4451.12 | 167.84 | 2133.42 | 599.68 | 1.06 | 0.03612754 | 4.89511967 | 1.09 | 1.99 | 1 |
| Sele | 28.76 | 4.66 | 19.1 | 1.31 | 0.59 | 0.03617607 | 3.95026588 | 0.96 | 1.23 | 1 |
| Twist1 | 151.58 | 74.56 | 42.4 | 21.21 | 1.84 | 0.03639847 | 3.12655973 | 0.98 | 3.71 | 1 |
| Clec7a | 193.94 | 100.21 | 52.86 | 14.12 | 1.88 | 0.03648508 | 3.77973127 | 0.84 | 4.01 | 1 |
| Eno1 | 31307.47 | 2421.12 | 47870.11 | 8498.43 | -0.61 | 0.03679528 | -3.83445191 | 0.41 | 0.07 | 1 |
| Ikbkg | 429.49 | 37.9 | 346.18 | 14.61 | 0.31 | 0.03685031 | 3.74982476 | 0.97 | 0.67 | 1 |
| Ly9 | 53.55 | 13.66 | 29.42 | 4.77 | 0.86 | 0.03706447 | 3.35423756 | 1.03 | 1.68 | 1 |
| Cd38 | 270.7 | 68.31 | 150.88 | 33.61 | 0.84 | 0.03715551 | 3.09155726 | 0.98 | 1.71 | 1 |
| Apoe | 9525.14 | 555.92 | 7125.01 | 902.7 | 0.42 | 0.03728408 | 3.70922089 | 0.95 | 0.9 | 1 |
| Cxcl12 | 147.14 | 49.33 | 64.93 | 17.33 | 1.18 | 0.03750589 | 3.25157905 | 1.02 | 2.34 | 1 |
| Pold1 | 255.75 | 22.42 | 342.87 | 43.76 | -0.42 | 0.03758243 | -3.24919772 | 0.56 | -0.01 | 1 |
| H2-DMa | 228.2 | 195.92 | 25.58 | 13.73 | 3.16 | 0.03771599 | 3.6487689 | 0.68 | 6.88 | 1 |
| Ccl11 | 432.34 | 225.83 | 125.43 | 48.99 | 1.79 | 0.03805893 | 3.31255388 | 1.05 | 3.5 | 1 |
| G6pdx | 1581.3 | 395.66 | 763.28 | 32.58 | 1.05 | 0.03820043 | 4.70962524 | 1.06 | 2.01 | 1 |
| Ms4a6b | 251.06 | 116.47 | 75.53 | 40.7 | 1.73 | 0.03837121 | 3.05661821 | 0.95 | 3.54 | 1 |
| Atm | 232.93 | 74.22 | 102.45 | 33.08 | 1.19 | 0.03853266 | 3.04429507 | 0.96 | 2.42 | 1 |
| Il18r1 | 42.62 | 14.29 | 19.1 | 1.31 | 1.16 | 0.03855963 | 4.48906231 | 1.03 | 2.27 | 1 |
| Arg1 | 99.57 | 17.4 | 33.87 | 15.2 | 1.56 | 0.04049299 | 3.79217625 | 0.86 | 3.32 | 1 |
| Fcgr1 | 369.71 | 209.36 | 85.64 | 21.31 | 2.11 | 0.0416584 | 3.82217836 | 0.83 | 4.49 | 1 |
| Herc6 | 1783.66 | 1060.32 | 357.26 | 59.11 | 2.32 | 0.04216298 | 4.17243004 | 0.95 | 4.71 | 1 |
| Ccl6 | 235.01 | 103.08 | 85.37 | 25.98 | 1.46 | 0.04222108 | 3.09039068 | 0.97 | 2.97 | 1 |
| Mtor | 246.34 | 1.2 | 223.75 | 8.15 | 0.14 | 0.04259421 | 4.52833509 | 1 | 0.27 | 1 |
| Eif2ak2 | 865.9 | 127.46 | 227.61 | 114.56 | 1.93 | 0.04386359 | 4.04981613 | 0.92 | 3.98 | 1 |
| Il18bp | 1018.05 | 702.76 | 138.75 | 8.16 | 2.88 | 0.04407858 | 4.54763269 | 1.11 | 5.6 | 1 |
| Irf8 | 1086.79 | 1297.5 | 40.73 | 3.55 | 4.74 | 0.04410873 | 4.55685568 | 1.2 | 9.21 | 1 |
| Thy1 | 493.54 | 187.68 | 205.23 | 37.06 | 1.27 | 0.04435548 | 3.54239154 | 0.83 | 2.8 | 1 |
| Ifitm1 | 70.06 | 33.24 | 19.1 | 1.31 | 1.88 | 0.0444495 | 4.42520618 | 1.04 | 3.7 | 1 |
| Apol6 | 332.58 | 340.39 | 19.1 | 1.31 | 4.12 | 0.04445124 | 4.54770994 | 1.17 | 8.02 | 1 |
| Ccl21a | 448.99 | 258.14 | 151.08 | 50.75 | 1.57 | 0.04591265 | 3.1522944 | 0.99 | 3.16 | 1 |
| Il2 | 23.41 | 2.23 | 19.1 | 1.31 | 0.29 | 0.04640641 | 2.98601532 | 0.99 | 0.61 | 1 |
| Met | 13596.98 | 1023.23 | 17129.79 | 1826.5 | -0.33 | 0.04641107 | -3.00668812 | 0.62 | 0.02 | 1 |
| P4ha1 | 1095.29 | 95.76 | 717.94 | 142.2 | 0.61 | 0.04652752 | 3.46032596 | 0.9 | 1.37 | 1 |
| Bid | 485.88 | 134.39 | 238.65 | 14.25 | 1.03 | 0.04652786 | 4.15106964 | 0.97 | 2.09 | 1 |
| Wnt4 | 41.8 | 4.6 | 24 | 5.74 | 0.8 | 0.0468772 | 3.49388504 | 0.88 | 1.79 | 1 |
| Cd27 | 23.68 | 2.39 | 19.1 | 1.31 | 0.31 | 0.04742223 | 3.01452446 | 0.99 | 0.64 | 1 |
| Mb21d1 | 73.47 | 14.05 | 48.44 | 4.93 | 0.6 | 0.04892237 | 3.23643804 | 0.87 | 1.4 | 1 |
| Gimap4 | 191.58 | 47.72 | 106.05 | 30.31 | 0.85 | 0.04975891 | 2.82166672 | 0.93 | 1.82 | 1 |
| C2 | 2314.98 | 1068.21 | 676.12 | 12.62 | 1.78 | 0.04988053 | 4.2966609 | 1 | 3.55 | 1 |
